# Why sensory neurons are tuned to multiple stimulus features

**DOI:** 10.1101/2020.12.29.424235

**Authors:** Matthew V. Macellaio, Bing Liu, Jeffrey M. Beck, Leslie C. Osborne

## Abstract

Many sensory neurons encode information about more than one stimulus feature. Multidimensional tuning increases ambiguity in stimulus-response relationships, but we find that it also offers an unexpected computational advantage, allowing the brain to better reconstruct sensory stimuli. From the responses of sensory neurons, populations, and sensory-driven movement behavior, more information can be recovered about a stimulus vector than about its individual components. We term this coding advantage “stimulus synergy” and show that it is distinct from other coding synergies, arising from inseparability of the response-conditioned stimulus distribution along individual stimulus dimensions. From extracellular recordings in motion sensitive cortex and measurements of pursuit eye movements, we demonstrate that stimulus synergy in cortical populations is preserved downstream in the precision of pursuit, and that a common decoding model predicts the level of synergy in pursuit behavior. This suggests that the brain exploits the information advantage afforded by multidimensional sensory tuning.

---

**S**ensory signals are multiplexed throughout the brain such that spike trains encode information about more than one stimulus feature: sound frequency and intensity, color and orientation, odor identity and concentration, body acceleration and retinal motion, etc. Sensory multiplexing might arise naturally from correlations in the environment, yet even nominally independent stimulus features are jointly represented by sensory neurons. Do multidimensional representations arise solely from anatomic or metabolic necessity, or is there a computational advantage? At first blush, representing more than one stimulus feature would appear to degrade rather than enhance sensory coding. When two or more stimulus features (which we refer to as dimensions) modulate a neuron’s responses, stimulus-response relationships become more ambiguous at the unit level, and increasing uncertainty about stimulus identity at the population level unless the population is perfectly homogeneous^1–3^. The problem is particularly acute in low signal to noise conditions or when rapid estimates are required such that spike counts are low and population activity is sparse^1,4,5^. In fact, we find that multidimensional tuning greatly constrains decoding operations, requiring the most efficient information recovery in order to explain behavior. Why then are multidimensional sensory maps so ubiquitous in the brain?

Many factors will contribute to the total amount of information encoded by a sensory population. Previous studies of multidimensional coding have largely focused on discriminability along a single stimulus dimension and how errors depend on the sharpness of feature tuning^1,2,6^, population size and response overlap^1,2,6,7^, heterogeneity in selectivity^1,2^, and the structure of correlations between neurons^1,8,9^. Implicit in single-feature discriminability analysis is the supposition that the brain estimates each stimulus dimension separately, or at least separably, but there is little evidence for or against this hypothesis. In the natural world the brain may need to derive veridical estimates of multiple stimulus dimensions simultaneously in order to guide appropriate behavior. If we are to understand how the brain derives sensory information from internal activity, we need to consider sensory estimation in a larger context – in the natural dimensionality of the representation.

We find that multidimensional representations offer an unexpected computational advantage: they encode more information about the collection of stimulus features than the total amount of information they encode about each stimulus feature independently – even when those stimulus dimensions and their representation are a priori independent. We term the information advantage “stimulus synergy”, drawing an analogy to coding synergies wherein a pattern of responses, like spike times within a train or cell activation within a group, collectively encodes more information than the sum of its component parts^10–13^. Stimulus synergy, however, arises from multidimensional nature of feature tuning rather than from patterning in neural responses. This has implications for downstream processing of sensory activity and sensory-guided behavior. Rather than estimate each stimulus dimension separately, we show that the brain could recover more information from individual neurons and populations of neurons by estimating all stimulus dimensions collectively. We also present evidence that the brain does in fact exploit the synergy afforded in multidimensional sensory representations through analysis of a movement behavior tightly coupled to sensory estimates. In fact, we show that behavior is only explained by a maximally efficient decoder that recovers synergy from the sensory population.

This study exploits the close connection between visual motion estimates and smooth pursuit behavior to examine how brain interprets multidimensional sensory signals. In the primate brain, visual motion is represented in the activity of cortical neurons in the extrastriate middle temporal area (MT), many of which respond selectively to retinal image motion and are tuned to varying degrees for multiple motion features such as direction, speed, binocular disparity, and spatial frequency^14–16^. MT mediates both the perception of motion^17^ and motion-driven motor behaviors like smooth pursuit eye movements^18^. Analysis of eye movement variation in pursuit eye movements^19–21^, its comparability to motion perception^21,22^, and cortical and cerebellar spiking^23,24^ suggests that little noise is added in downstream motor processing. Under favorable conditions, the initial eye movement can report the internal estimate of target direction and speed recovered from cortical activity with surprising fidelity. Using a combination of extracellular recording, eye movement measurements and simulations, we explore the implications of a multidimensional sensory code for motion on information recovery about direction and speed at the level of individual neurons, populations and in sensory-driven movement behavior.

## Results

When a sensory neuron’s firing rate is modulated by multiple features of a sensory stimulus, the same firing rate can correspond to many combinations of those features, increasing ambiguity in the neural representation that must be disambiguated by downstream circuits. The number of stimulus feature combinations that can give rise to the same firing rate increases with the number of encoded features, which we will term dimensions (D). Clearly, the number of possible stimulus values that correspond to the same neural response increases with the dimensionality of the tuning function from two points in 1D, to a ring in 2D, to a spherical shell in 3D, etc. (Figure 1A). We presume that the brain resolves the ambiguity by interpreting the activity of many such neurons with a range of feature selectivities, but how? One possibility is that the brain decodes estimates of each stimulus dimension separately (Figure 1C); another is that the decoder operates in a higher dimensionality perhaps even that of the representation so that it can estimate the multidimensional (nD) stimulus as a whole (Figure 1B). These two strategies are not computationally identical and can lead to different estimates of the stimulus. Consider a simple example where a model neuron’s firing rate is independently modulated by 2 stimulus dimensions with identical Gaussian tuning functions (Figure 1E, left panels). The maximal firing rate is elicited by a single stimulus vector 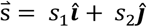, but many combinations of *s*_1_ and *s*_2_ can give rise to the same intermediate response value. To an observer who considered only s_1_ to be of interest, a count of *n*, if less than the maximal count, would indicate that s_1_ likely had one of 2 values corresponding to the intersections of the red dashed line with the tuning function (Figure 1E, top left panel). However, in practice, a count *n* could arise from any combination of s_1_ and s_2_ values lying on a circle (Figure 1D, red dashed circle) defined by the outer product of the 1D tuning functions. Spiking variability increases uncertainty about the stimulus, such that the probability of stimulus values, 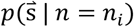, broadens from a circle (Figure 1D) to a torus (Figure 1E, right panel), though the stimuli corresponding to the highest likelihoods do not change. Separate estimates of each stimulus dimension, s_1_ or s_2_, are potentially significantly less certain and less accurate than a joint estimate of the stimulus vector. The likelihood of observing a value of s_1_ (or s_2_) given a count of *n_i_* is a bimodal distribution whose peaks correspond to the intersection points in the left panels of Figure 1E (Figure 1F, left panels). Combining the likelihoods of s_1_ and s_2_ yields the 2D function shown in the right-hand panel, representing the outer product of the bimodal distributions (Figure 1F). The color density is more broadly distributed in Figure 1F than in 1E, indicating increased uncertainty about the identity of the stimulus. But not only is the total stimulus less well specified, the maximum likelihood estimates of its parameters are also incorrect. The likeliest stimulus estimates correspond to the dark red peaks in Figure 1F (right panel), which correspond to the peaks of the outer product of the marginal distributions on the left, and those values do not lie on the circle defining the true expected values of s_1_ and s_2_ (white dashed circle, Figure 1F). The differences between Figures 1E and 1F suggest that a multi-dimensionally tuned neuron can specify the stimulus vector more precisely than it specifies its individual dimensions, i.e. that

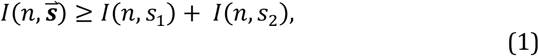

where *I()* terms represent mutual information. We term this extra information, when positive, “stimulus synergy”. The term synergy implies that the whole -- the total encoded information about the multidimensional stimulus -- is more than the sum of its component parts. The fact that a Poisson model displays synergy indicates that the coding advantage does not arise from correlations between spike times that allow a pattern of spikes to encode more stimulus information that the total count^10^. Nor does the model’s synergy arise from nonlinear interactions between stimulus features, such as a “face cell” that fires in response to a face but not to its component feature presented individually or in the wrong spatial arrangement^25^. All were absent by design. Rather, stimulus synergy appears to be inherent to multidimensional representation. Indeed, we will show in a later section that if two different stimulus dimensions are a priori independent then stimulus synergy cannot be negative and thus coding cannot be redundant. As a result, most sensory neurons will display some degree of stimulus synergy, with potential implications for how downstream targets should decode those spikes most efficiently. We tested this hypothesis by recording from cortical neurons that respond to multiple independent features of visual motion and measuring the information encoded about the multidimensional stimulus vector compared to its components.

**Figure 1.**
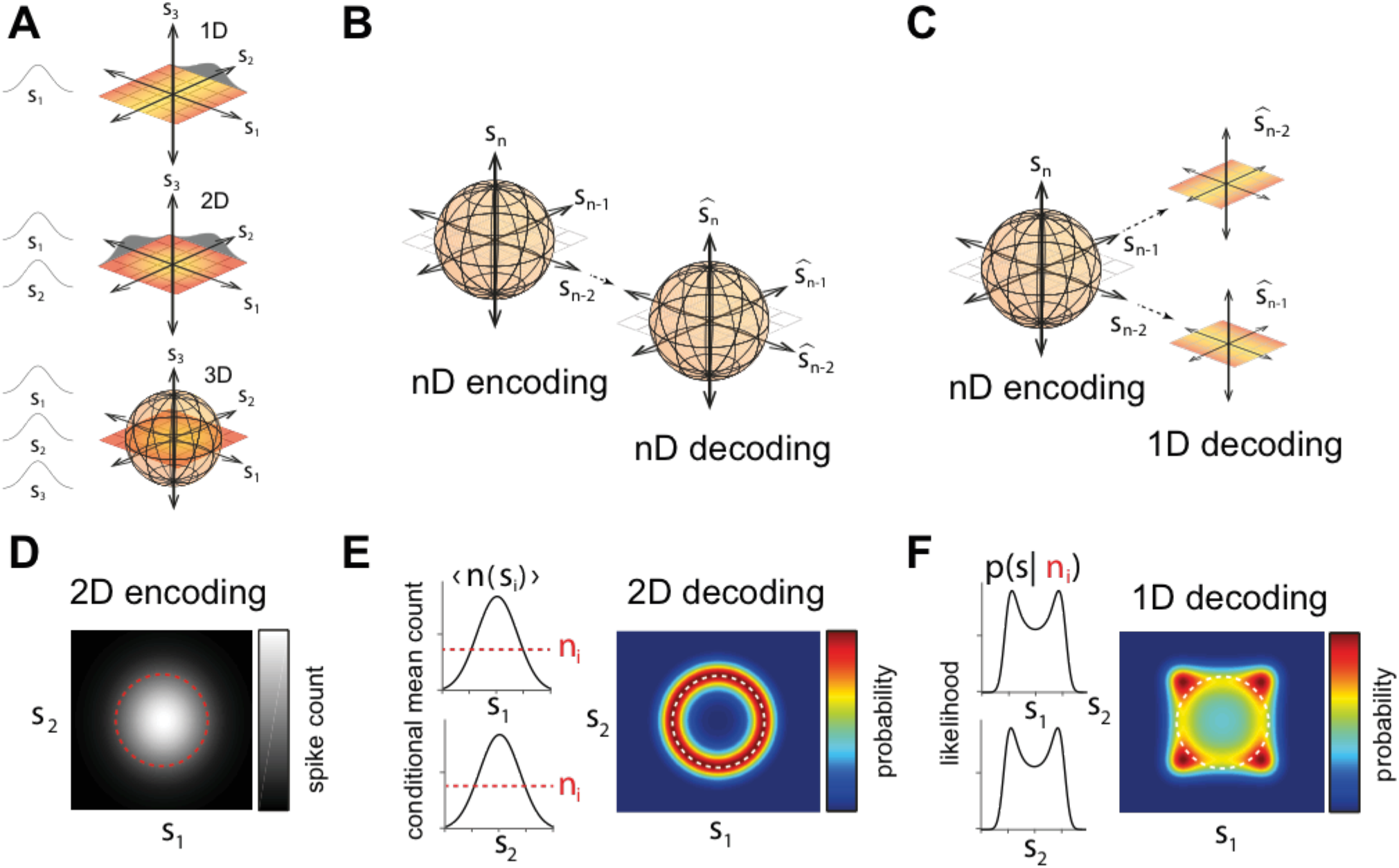
Multidimensional tuning and stimulus specificity. **A.** Illustration of dimensionality and encoding. Color gradients indicate a modulation in response with stimulus value (s_1_, s_2_, s_3_, … s_n_), also indicated by the distributions at margins. The responses of a neuron tuned to *n* stimulus dimensions can be read out with respect to all dimensions together (**B**) or each dimension individually (**C**). **D**. Illustration of the 2D stimulus-conditioned mean count (right) characterized by identical, Gaussian tuned responses to each stimulus dimension (left). Red dashed lines (left) and white dashed circle (right) correspond to an average count value of *n_i_*. Ambiguity in stimulus identification depends on the dimensionality of the representation. **E**. Conditional 2D stimulus distribution for a Poisson model with the tuning functions in (D). The most likely values of s coincide with the dashed circle (re-plotted from D) **F**. If each stimulus dimension were estimated independently, the 1D conditional stimulus distributions are bimodal (left panels). The outer product of those 1D conditional distributions (right panel) describes the uncertainty about the vector stimulus. Note that neither the peaks of the 2D distribution (dark red color) nor the mean lie on the white circle indicating the true stimulus values. The greater extent of the color shading in F vs. E indicates that the total uncertainty is greater if stimulus dimensions are estimated independently rather than jointly.

### MT neurons encode motion direction and speed synergistically

While the monkeys fixated, we recorded extracellular activity from isolated MT units responding to 200 ms - steps of coherent random dot patterns moving at a fixed direction and speed (Figure 2A; see Methods). Ninety-six motion stimuli (12 directions, 8 speeds) were repeated an average of 33 times, which allowed us to compute the terms in Equation 1 directly from the recorded spike counts.

**Figure 2.**
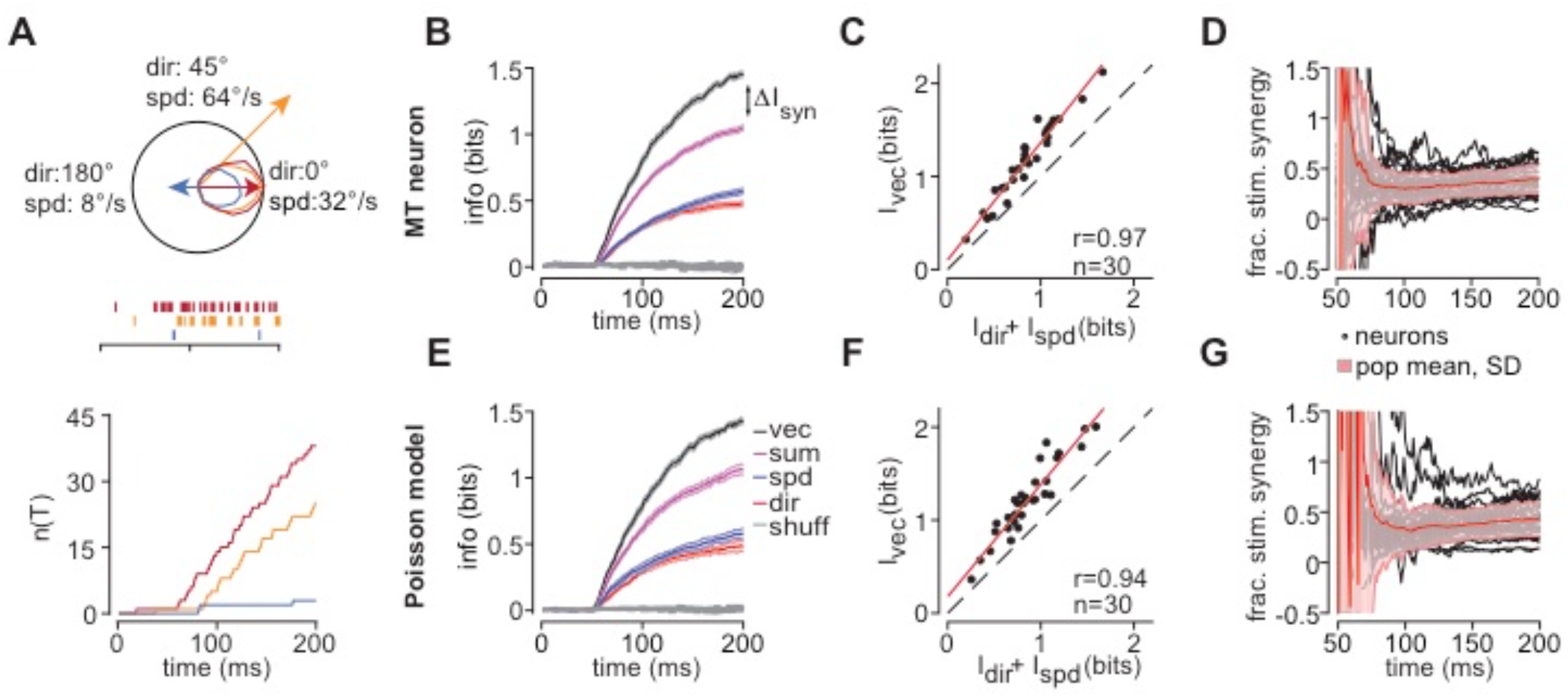
MT neurons encode stimulus components synergistically even when temporal spike patterns are disrupted. **A.** We used cumulative spike count to quantify MT responses over time, shown for 3 directions and speeds for an example unit (see tuning curves, upper panel). Spike times (colored dashes, middle panel) were binned at 1 ms. Cumulative spike counts (lower panel) were computed from motion onset 0 ms). **B.** Mutual information between spike count and stimulus over time for an example unit. Mean and errorbars (SD) have been corrected for sample size (see Methods). Information about the 2D motion vector (black) is greater than the sum (magenta) of the 1D information about direction (red) and speed (blue). Gray trace indicates the same calculation applied to stimulus-label-shuffled data as a control. **C**. Population data from 30 units (dots) sampled at 200 ms relative to motion onset. Most neurons encode stimulus dimensions synergistically. A linear fit (red) to the data has a slope greater than unity (dashed) (I_vec_ = 1.26*I_sum_). **D.** Stimulus synergy as a fraction of summed direction and speed information for all units (gray dotted lines), and population mean, SD (red shading) is approximately stable over time after response onset of neurons (~65 ms from motion onset). **E, F, G**. Same analyses applied to Poisson models of each unit, created by randomizing trial identity at each time step to break within-trial temporal correlations without altering the PSTH (see text). The randomization creates Poisson-like spiking statistics but preserves the features of stimulus synergy in the MT sample. Stimulus synergy therefore does not arise from temporal patterning of spike times.

For each unit, we compared information encoded about the 2D motion vector, 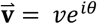, to the information encoded about direction (*θ*) and speed (*v*) individually. We measured the cumulative spike count, *n*(*T*), at 1ms intervals from motion onset. Spike counts increased quickly for preferred stimuli (red), more slowly for nonpreferred stimuli (gold), and spiking was suppressed for directions in the anti-preferred direction (blue) (Figure 2A). We computed the mutual information between *n*(*T*) and motion direction *I*_dir_(*T*) = *I*(*n*(*T*), *θ*), speed *I*_spd_(*T*) = *I*(*n*(*T*), *v*), and the 2D motion vector 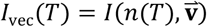, at each time step (see Methods,). Figure 2B shows the information time courses for each from an example MT unit. Traces represent the extrapolated information values at infinite sample size, and the errorbars are extrapolated standard deviations^26^. Information about motion speed (blue) and direction (red) both increase quickly after response onset and thereafter more slowly^27^. After 200 ms of stimulus motion, the example unit in Figure 2B encodes 38.1% (0.40 bits) more information about the stimulus motion vector (black trace) than the summed information about direction and speed (magenta trace, Figure 2B), which was near the sample mean of 39.6±14.2% (SD, n=30 units; p =2.3e-14, t=-13.5, different from zero, 1-tailed t-test, Figure 2C). The level of stimulus synergy was stable throughout the response period (Figure 2D).

Previous studies have reported that synergy can arise from temporal patterning within spike trains^10^ and from nonlinear feature selectivity^28^. We confirmed that neither are the origin of stimulus synergy in MT neurons. We eliminated a potential contribution from temporal patterning by randomly shuffling the trial identity of each spike without changing its timing. Trial shuffling creates an inhomogeneous Poisson-like model of each unit by breaking temporal correlations between spikes without changing the time-varying firing rate^27^. Shuffling did not alter the level or the dynamics of stimulus synergy (Figures 2E-G). The model population had an average of 43.7 ± 17.9% (SD, n=30 model units; p=1.4e-13, t=-12.6, 1-tailed t-test) stimulus synergy after 200 ms of stimulus motion, which was not statistically different from the neural data (p=0.33, t=-0.98, n=30; two tailed 2 sample t-test). While temporal relationships between spikes can create a synergistic sensory code, they are not necessary for stimulus synergy.

We also ruled out nonlinear feature selectivity as the source of stimulus synergy in MT. Neurons with highly nonlinear selectivities might only respond to a particular conjunction of stimulus features (e.g. face-selective neurons). Such neurons are expected to display large amounts of stimulus synergy because they encode little to no information about individual stimulus features comprising the stimulus vector. Synergy in a neuron with separable tuning is not expected. If MT motion selectivities were perfectly separable then their 2D tuning functions would equal the outer product of their direction and speed tuning functions, *r*(*θ, v*) = *r*(*θ*)*r*(*v*). Deviations from separability would imply that direction tuning depended on speed and vice versa. We determined the degree to which the 2D trial-averaged direction-speed tuning functions were separable using singular value decomposition. We defined an index of separability (SI) as the fraction of variance captured by the largest singular value with a maximum value of 1 for perfect separability^29^ (see Methods). We found that MT tuning functions in our data sample deviated from perfect separability by modest amounts. The distribution of SI values measured 200ms after motion onset had a sample mean of 0.98 ± 0.01 (SD, n=30 units). For reference, taking the outer product of each unit’s direction and speed tuning functions to create a purely separable 2D tuning function (SI=1) and then rotating by 45° to maximize tuning nonlinearity reduces the SI to 0.88±0.05. The small deviations from pure separability in MT units were not significantly correlated with their fractional synergy levels (R = −0.04, p=0.82; one-tailed t-test). While inseparable tuning functions could certainly enhance stimulus synergy, it is neither necessary nor the origin of stimulus synergy in MT.

### Pursuit eye movements reflect the stimulus synergy observed in MT

Synergistic coding of stimulus features, in MT or elsewhere in the brain, has potential consequences for behavior. We tested for evidence that the brain exploits stimulus synergy by analyzing monkeys’ pursuit eye movements. To stabilize the retinal image of a moving target, the pursuit system translates MT activity into commands to the extra-ocular muscles to smoothly rotate the eye along with a moving target. Under appropriate conditions, the translation of visual activity to eye movement is very precise^19–21^ such that the eye movement reports the estimate of target motion formed during the time period when there is substantial retinal image motion because the eye has not yet caught up to the target. Monkeys tracked patterns of dots that translated across the screen with directions of 0°, ±10°, ±20°, ±30°, leftward and rightward at speeds of 10, 15, 20, 25, and 30°/s (Figure 3B). MT encodes motion speeds that are faster than the eye is capable of tracking, so we selected a range of directions and speeds that were comfortable for the monkeys to track. We confirmed that matching the direction range to the MT experiments did not impact results. We randomized presentation order to minimize anticipation.

**Figure 3.**
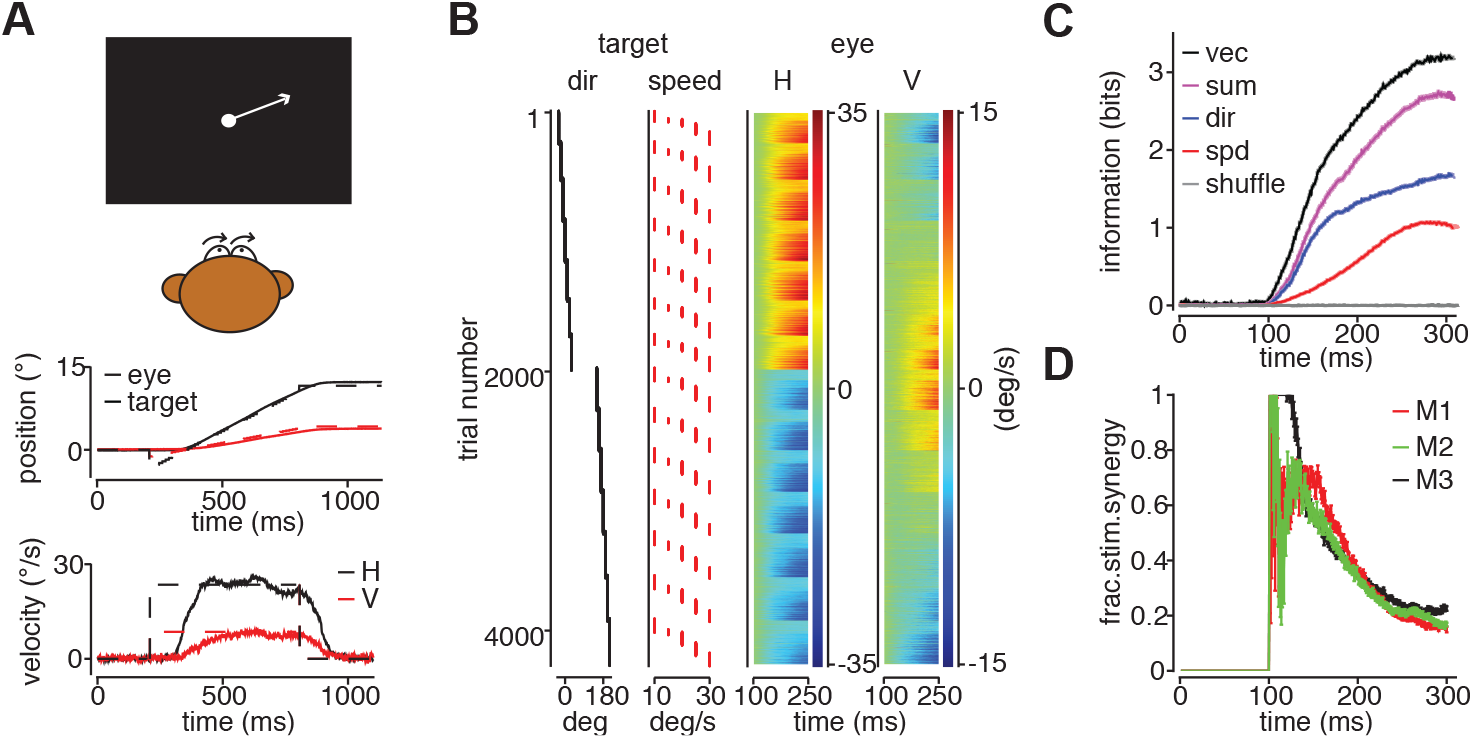
Stimulus synergy is apparent in pursuit behavior. **A.** (top panel) Cartoon of a pursuit experiment. Monkeys pursued spot targets that appeared eccentrically as the fixation point was extinguished then immediately began to translate toward the fixation position, with directions of 0°, ±10°, ±20°, ±30°, leftward and rightward at speeds of 10, 15, 20, 25, and 30°/s. Horizontal (H) and vertical (V) target (middle) and eye velocities (bottom) over time for a single pursuit trial with a target moving at 20° target moving at 25°/s. **B**. (Left panels) Target directions and speeds for a single experiment. (Right panels) Each row of the color density plot represents the horizontal (H, left panel) and vertical (V, right panel) eye velocity on a single trial during the first 125 ms of pursuit from a day’s experiment with M1. **C.** Mutual information between H, V eye velocity and target motion over time for M1. Information about the 2D motion vector (black) is greater than the sum of the information about direction (red) and speed (blue) individually. Times are relative to target motion onset. **D**. Fractional stimulus synergy decreases over time from pursuit initiation to ~0.2. Data from 3 monkeys.

Pursuit behavior shows substantial levels of stimulus synergy. Using data sets like the example in Figure 3B, we computed the mutual information between eye and target velocity components to determine if stimulus synergy in motion coding is reflected in eye movement behavior. To avoid a 1 bit jump in information at pursuit onset, we combined rightward and leftward target conditions with the same eccentricity relative to horizontal. At each time point, we discretized velocity values into bins and then computed mutual information, correcting for finite sample size as with the neural recordings. As with MT responses, the information about direction (red trace, Figure 3C) and speed (blue trace, Figure 3C) rises quickly after pursuit onset, ~100 ms after the target begins to move. Pursuit reaches a stable level of precision within ~200ms after the onset of eye movement such that information does not continue to increase^20^. The overall level of behavioral information remains substantially lower than the 5.12 bits of the stimulus set, but greater than information encoded by individual MT neurons, confirming intuition that the brain derives motion estimates from population responses^27^. Like MT neurons, more information is encoded about the target motion vector in pursuit behavior (black trace, Figure 3C), than the sum of the information about direction and speed (magenta trace, Figure 3C). Synergy levels were consistent among monkeys. Measured 250 ms after target motion onset, the fractional synergies for each animal are 24±0.9% (M1), 19±0.6% (M2), and 21±1.0% (M3) (SD, n=30 draws from 70% of total data sample) (Figure 3D). The pursuit results suggest that the brain may be able to exploit stimulus synergy to improve behavioral performance. To determine how that is possible, we analyzed stimulus synergy in the population representation of motion direction and speed and how that in turn impacts decoding multiple stimulus features.

### Stimulus synergy is preserved in MT population responses

The appearance of stimulus synergy in behavior implies that either the sensory population response retains some of the synergy in individual neural responses and the brain’s downstream circuity recovers that information, or that inefficient decoding creates or enhances behavioral synergy from a relatively non-synergistic neural population code. In the context of current theories of sensory decoding, it is not immediately obvious which of these possibilities is the case. Ambiguity about stimulus direction and speed in the response of a single unit should be greatly reduced in the response of a large, diversely tuned population. For example, the “donut” of uncertainty about stimulus values in the model unit shown in Figure 1E, could shrink to a small point of intersection when the conditional stimulus profiles of many units overlap to create an unambiguous and thus non-synergistic code for stimulus identity. However, factors such as noise correlations between neurons and heterogeneity in the distribution of tuning parameters that limit the reduction in ambiguity could create the potential for stimulus synergy in the population response.

We analyzed stimulus synergy in MT population responses and tested several decoding models to determine if the extra information from stimulus synergy is recoverable downstream. To create realistically diverse population responses, we randomly selected preferred directions and speeds, tuning bandwidths, spontaneous and peak firing rates from distributions that were consistent with our recorded data and the literature (details in Methods; Fig. 4A, B). We generated conditional response covariance matrices to reflect the presence of baseline correlations, correlations that decayed as a function of the difference between preferred stimuli, and “information limiting” correlations^30–32^. Due to inhomogeneities in our populations, the information capacity of our network scaled with the number of neurons^9^. We therefore added differential correlations^32^ so that total information eventually saturates to a level that represents the total amount of input information. Simulated neural correlations did not mimic direction and speed correlations during stimulus presentation and thus did not spuriously create synergy. Rather, we instantiated a priori independent differential correlations by effectively adding independent noise to the presented speed and direction. This is equivalent to assuming that the stimulus that drives the population is given by the true value of the stimulus plus some noise. We scaled neural correlations such that the maximum Fisher information of the population response exceeded that in motion estimation behaviors. Motion discrimination thresholds, a measure of sensory information, are 3° for direction and 10% for speed in both perception and pursuit^21^.

**Figure 4.**
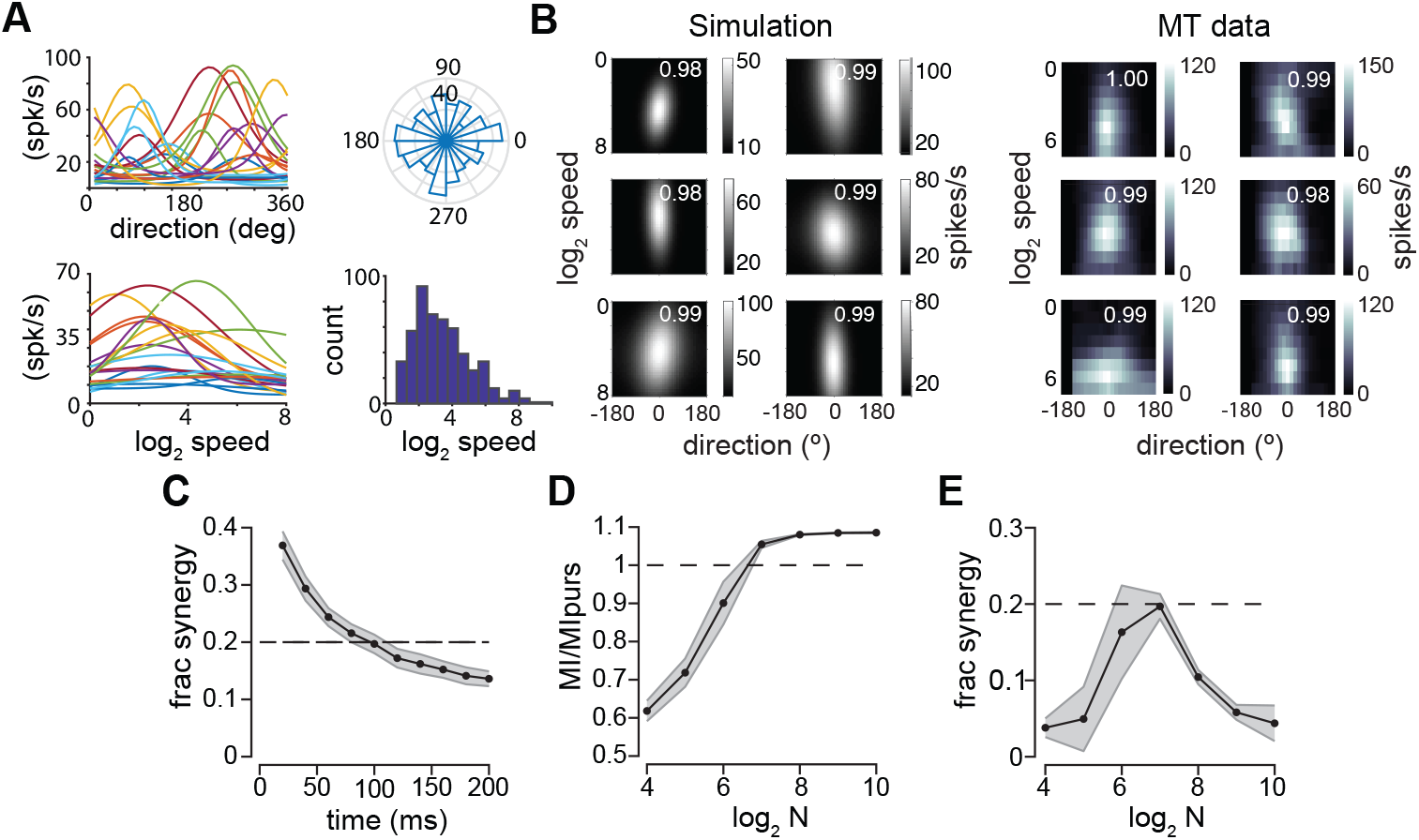
Stimulus synergy is preserved in a diverse sensory population. **A** Direction-speed tuning functions for example MT units. Left panels, direction (top) and speed (bottom) tuning functions for randomly selected, simulated units. Right panels, distribution of preferred directions (top) and speeds (bottom) for 256 simulated units match published statistics. **B.** (Left panels) 2D conditional mean count for 6 simulated units. Separability index (SI) values, upper right, matched to the distribution in our data sample. (Right panels) same for 6 MT units. **C**. Fractional stimulus synergy in the population response over time. Values represent mean over 40 randomly generated 128-unit populations, errorbars SD. Dashed black line represents output information observed in behavior. **D**. Population MI with respect to output information estimated from the precision of pursuit, plotted vs. log_2_ population size. Values are mean over 40 simulations, error bars are SD. **E**. Fractional synergy values vs log_2_ population size for simulations in D.

Our simulation approach allowed us to analyze the information capacity and synergy of the population analytically, without generating samples of activity. This was accomplished by first computing linear Fisher information (FI), a quantity that depends solely on the gradient of the conditional mean population response and the stimulus conditioned noise covariance. For a vector stimulus, FI is a matrix whose diagonal values reflect signal-to-noise (SNR) in population activity about each stimulus dimension, while the off-diagonal elements reflect the interactions between dimensions (Eqn. 13, Methods). We applied the Brunel and Nadal approximation^1^ to convert Fisher information matrices into mutual information in bits. We found that MT populations display a substantial amount of stimulus synergy, comparable to the level observed in pursuit behavior (Figure 4C). Firing rates remained constant in our simulations, so the dynamics in information quantities and fractional synergy in Figure 4C are due only to accumulating signal to noise with ongoing firing. We report population information values with respect to the level of motion vector information observed in pursuit, computed from behavioral direction and speed precision. In most of our simulations we assumed that input (Fisher) information (i.e. the limit on population performance) exceeded behavior by a factor of 2. Using a factor of 2, the output information of realistically diverse MT populations only slightly exceeded behavioral information levels. Increasing that factor raised both the level of output information and synergy.

Information and synergy levels increased with increasing population size, such that both peak for populations of 128 cells (Figure 4D, E). Near this peak, adding more neurons to the coding pool would have little behavioral benefit relative to the metabolic cost of adding more neurons or spikes. The fact that synergy reliably peaks in the regime where information levels begin to saturate as a function of pool size suggests that observing substantial levels of synergy in cortex may be indicative of an energy efficient neural code. That is, the presence of significant amounts of synergy in MT and pursuit suggests that the neural code is being optimized to maximize behaviorally-relevant output information subject to some constraint on metabolic cost.

### Decoding population responses to predict pursuit information and synergy

It is important to note that the presence of synergy in MT does not necessarily lead to synergy in behavior-output synergy depends on the decoder. A decoder that can exploit the stimulus synergy available in the population response to improve stimulus estimation should be sensitive to the direction-speed interaction terms of the Fisher information matrix (see Eqn. 13). A decoder whose weights are derived from the full Fisher information matrix would be maximally efficient, likely to maximize information recovery as well as synergy. However, decoding inefficiencies could also create synergistic outputs. This is because decoders of neural activity can be biased or inefficient in different ways, potentially enhancing or reducing synergy in the decoded variables. Therefore, we searched for decoding mechanisms that could both recover sufficient information from MT while maintaining levels of synergy consistent with our behavioral results. We tested 4 previously proposed types of decoders that varied in their knowledge of, and ability to exploit, both synergy and noise covariance in the MT population. We compared 1D and 2D versions of a population vector decoder (PV), a global linear decoder (GL), a naïve Bayesian decoder (NB), and locally-optimal linear estimation (LOLE), our proxy for maximum-likelihood decoding (ML). All decoders are described in detail in the Methods, but we summarize the important differences here. The PV decoder weights each unit by its preferred stimulus. The GL decoder weights each unit by its response variance with respect to the stimulus-averaged noise covariance. The weights of the LOLE decoders are derived from the Fisher information matrix and therefore has full knowledge of the stimulus dependence of the noise covariance matrix. The NB decoder, on the other hand, only has knowledge of the diagonal elements of that matrix. We applied these decoders to our simulated population and computed the mutual information between the true value of the stimulus and the decoded value of the stimulus. The NB and LOLE decoders are unbiased by design, meaning that the decoder weights are selected to ensure that stimulus estimation errors average zero for each stimulus condition and information calculations are straightforward. The GL and PV decoders are biased, such that their direction estimates vary with stimulus speed and vice versa. To evaluate the information recovered by the biased decoders, we directly computed the bias and variance of their stimulus estimates, then converted these values to decoder-specific Fisher information from which we could estimate the mutual information (Brunel and Nadal 1998). For each decoder type, we also computed stimulus synergy as fractional difference between the information recovered by the 2D decoder and the summed information about direction and speed from their 1D counterparts. None of the decoders returned redundant stimulus estimates (synergy < 0) (Figure 5B).

**Figure 5.**
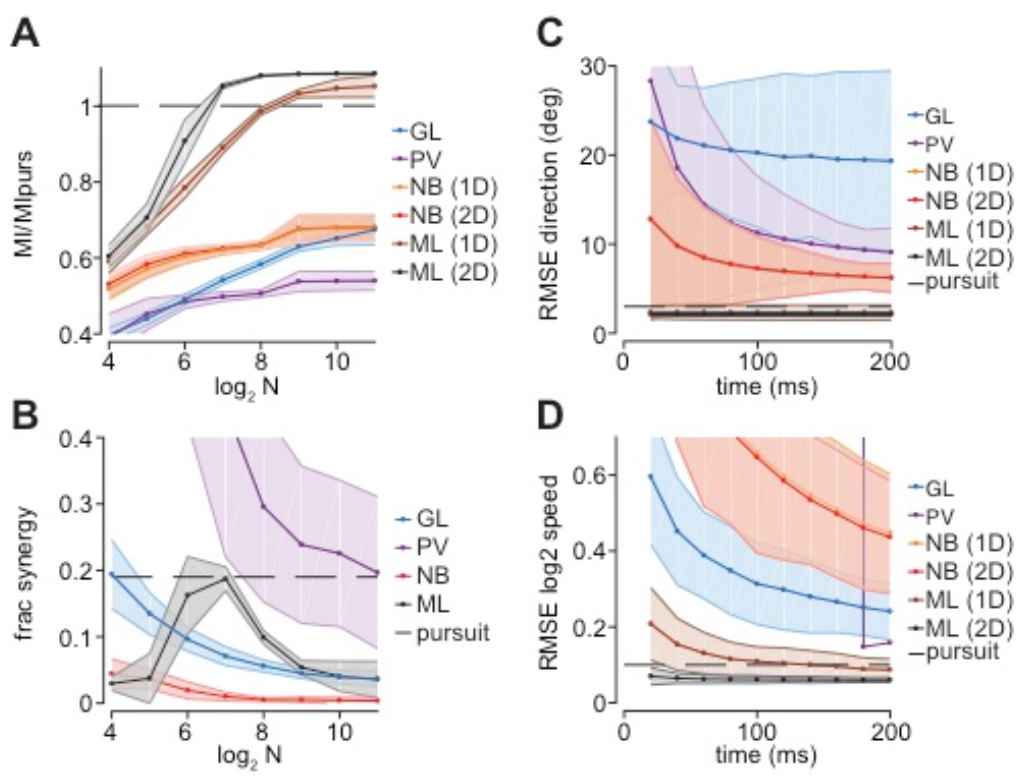
A 2D maximum likelihood decoder predicts behavioral information and synergy levels. **A.** Mutual information about the 2D stimulus vector with respect to log_2_ population size. Each trace represents a different decoder (see text). MI values were computed were computed from decoded stimulus estimates (variance, bias) except for ML 2D which allows direct calculation of Fisher information. Information values are reported as a fraction of output information estimated from the precision of pursuit. Curves represent mean values over 40 simulations, error bars SD. **B**. Fractional stimulus synergy as a function of log_2_ population size. For each decoder type (GL, PV, NB, ML) curves represent the ratio of information about the 2D motion vector from the 2D decoder to the summed information about direction and speed from its 1D counterparts. Dashed line indicates the degree of stimulus synergy measured in pursuit behavior. **C**. Errors in estimating stimulus direction and **D**. speed made by each decoder as a function of integration time. Only the 2D ML decoder (LOLE) exceeds the precision observed in pursuit (dashed black line). Values represent mean, SD over 40, 128-unit populations.

The maximum likelihood (2D LOLE) decoder out-performed all other decoders in both efficiency and output synergy (black traces, Figure 5). Both the PV (purple traces Figure 5A, B) and LOLE (black traces, Figure 5A, B) decoder had substantial levels of stimulus synergy, but only the LOLE was able to recover both the amount of motion information and the level of stimulus synergy reflected in pursuit behavior. Figures 5C, D show the scale of direction and speed errors made by each decoder. The 2D ML decoder had the smallest error levels (black traces, Figure 5C, D), but its direction-only and speed-only 1D counterparts (brown traces Figure 5C, D) also performed comparably to behavior. Neither the most commonly used decoders (GL and PV) nor the NB decoders recovered sufficient information to account for pursuit because they ignore features of the population response. The GL decoder conflates signal and noise correlations while the PV decoder fails to take into account variability in signal to noise ratios of individual units and how they change with the multidimensional stimulus. As a result, these decoders made direction and speed estimation errors that exceeded those in pursuit (purple, blue traces Figure 5C, D). For example, a population vector decoder produced direction errors that were ~4 times larger than pursuit, and speed estimates were unstable on a behaviorally relevant time scale for motion integration except for the largest populations (purple trace, Figure 5C, D). The NB decoder performed somewhat better (orange, red traces Figure 5) but not as well as the ML (LOLE) decoder.

Three decoders showed substantial output synergy (PV, GL, ML), but for different reasons. Figure 5B shows the difference in information about the stimulus vector compared to estimating direction and speed independently for each decoder. The ML decoder outputs are synergistic because it extracts the full information from the synergistic population response. The linear GL and nonlinear PV decoders show stimulus synergy due to their inefficiency (purple, blue traces Figure 5B). Although each treats direction and speed independently, the PV and GL decoders induce correlations between direction and speed estimates that in turn create stimulus synergy. Those correlations lower output information, creating poor stimulus estimation performance (blue, purple traces Figure 5 C, D). As such, neither the GL nor PV decoders could explain our behavioral data.

Of the tested decoders, only optimal vector-decoding of both speed and direction (2D-LOLE) generated levels of synergy and information comparable to behavior (ML-2D, black traces, Figure 5). Curiously, this level of synergy was only present for populations that were large enough that the input information, controlled by differential correlations^32^, was close to the observed level of output information, but populations not so large that input information recovery was almost perfect. In retrospect, this makes sense. The information capacity of heterogeneous networks with realistic correlation structures is known to scale with the number of neurons^9,32^. Therefore, in the presence of finite input information, output information necessarily saturates to input information levels as the number of neurons increases and synergy goes to zero. This result suggests that the presence of significant synergy in MT and pursuit likely results from a network structure that optimizes the recovery of a finite amount of input information under a cost constraint which becomes increasingly expensive in terms of information-gain per spike.

### Average MT tuning bandwidths maximize stimulus synergy

To expand beyond the specific case of motion coding in MT, we used simulations to dissect the impacts of tuning bandwidth, tuning dimensionality, separability, tuning function form, and stimulus correlations to stimulus synergy. We parametrically changed the tuning bandwidth and number of encoding dimensions in a - model unit and computed the mutual information between spike count and stimulus as well as the fractional stimulus synergy for each combination. Our model units had identical, separable, Gaussian tuning functions for each stimulus dimension, like the example depicted in Figure 1. As expected, we found that higher levels of information are encoded with narrow tuning functions, <30% of the stimulus range, but only for a single dimension (Figure 6A). Increasing the dimensionality of a neuron’s selectivity requires broader tuning functions to avoid a steep information roll-off. Models with tuning functions similar to our MT sample (gray and black traces, Figure 6A-C) are well-suited to encode 2-3 dimensions which is consistent with MT’s role in encoding motion in depth^33^. As with information, broader tuning is necessary to maintain stimulus synergy for more 4 or more dimensions, even when synergy is expressed as a fraction the total information for each condition (Figure 6B). The simulation results predict that MT neurons should encode 20-25% more information about motion vectors than about direction and speed separately, which is about what we observe (compare black gray traces Figure 6B to Figure 3D).

**Figure 6.**
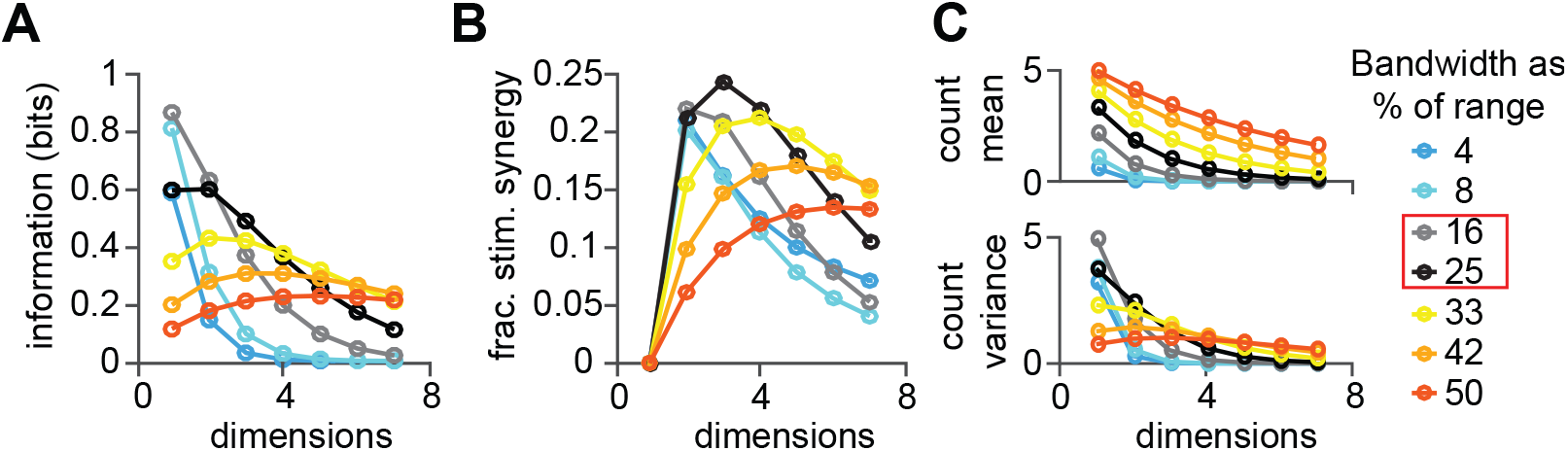
Optimal encoding dimensionality and level of stimulus synergy depend on stimulus tuning bandwidth. **A.** Mutual information between spike count and an arbitrary stimulus feature from a homogeneous Poisson model unit with identical response parameters for each stimulus dimension (see Methods for details). Model unit tuning bandwidth expressed as percentage of the tested range. Gray and black curves (red box) represent the range-normalized direction and speed bandwidths of the MT sample as a fraction of 360° (direction range) or 10 log_2_ units (speed range, 0.25-256°/s). **B**. Fractional stimulus synergy peaks at lower dimensionality at narrow tuning bandwidths (blue) than at wider bandwidths (orange). **C.** Mean spike count across stimuli increases with wider bandwidth at all dimensionalities (top panel). Variance of spike count across stimuli is greater at narrow bandwidths than wide bandwidths at low dimensionality, but this trend reverses at high dimensionality (bottom panel).

Less intuitive is why increasing the number of encoded dimensions impacts information and fractional stimulus synergy so differently based on tuning bandwidth. Adding coding dimensions decreases information and stimulus synergy in narrowly tuned model responses (blue curves, Figure 6A, B) but increases information and stimulus synergy for broadly tuned neurons (orange-red curves, Figure 6A, B). The reversal in the relationship between tuning sharpness and information or stimulus synergy can be explained by changes in the spike count distribution with the number of stimulus dimensions^34^. The number of stimulus values increases exponentially with the number of coding dimensions, but because the peak firing rate remains fixed, the maximum number of coding symbols remains the same (0 to the maximum count). Generally speaking, spreading the same number of coding symbols across a larger stimulus space lowers the average count (Figure 6C, top), and decreases count entropy and coding capacity. Particularly with narrow tuning functions, the effect of increasing stimulus dimensionality is a steep decline in count variance *across stimulus conditions* (blue curves, Figure 6C, bottom). However, the drop off in variance is more gradual for broadly tuned models, and even increases slightly from 1 dimension to 3 dimensions (red curves, Figure 6C, bottom). Broad tuning restricts the range of response values because all stimulus conditions elicit non-zero firing rates. However, with each added stimulus dimension, the minimum rate falls, increasing the response range (variance) and mitigating the tendency for the drop off in mean rate to lower the entropy. The amount of information encoded, and the degree of stimulus synergy reported in Figures 6A, B reflect the balance of the trends in Figure 6C. Tuning bandwidths in our MT sample correspond to models that maximize information and stimulus synergy for 2-3 dimensions. Non-identical tuning bandwidths between dimensions yield varying amounts of synergy, but do not materially change the trends in Figures 6A-B.

**Figure S2.**
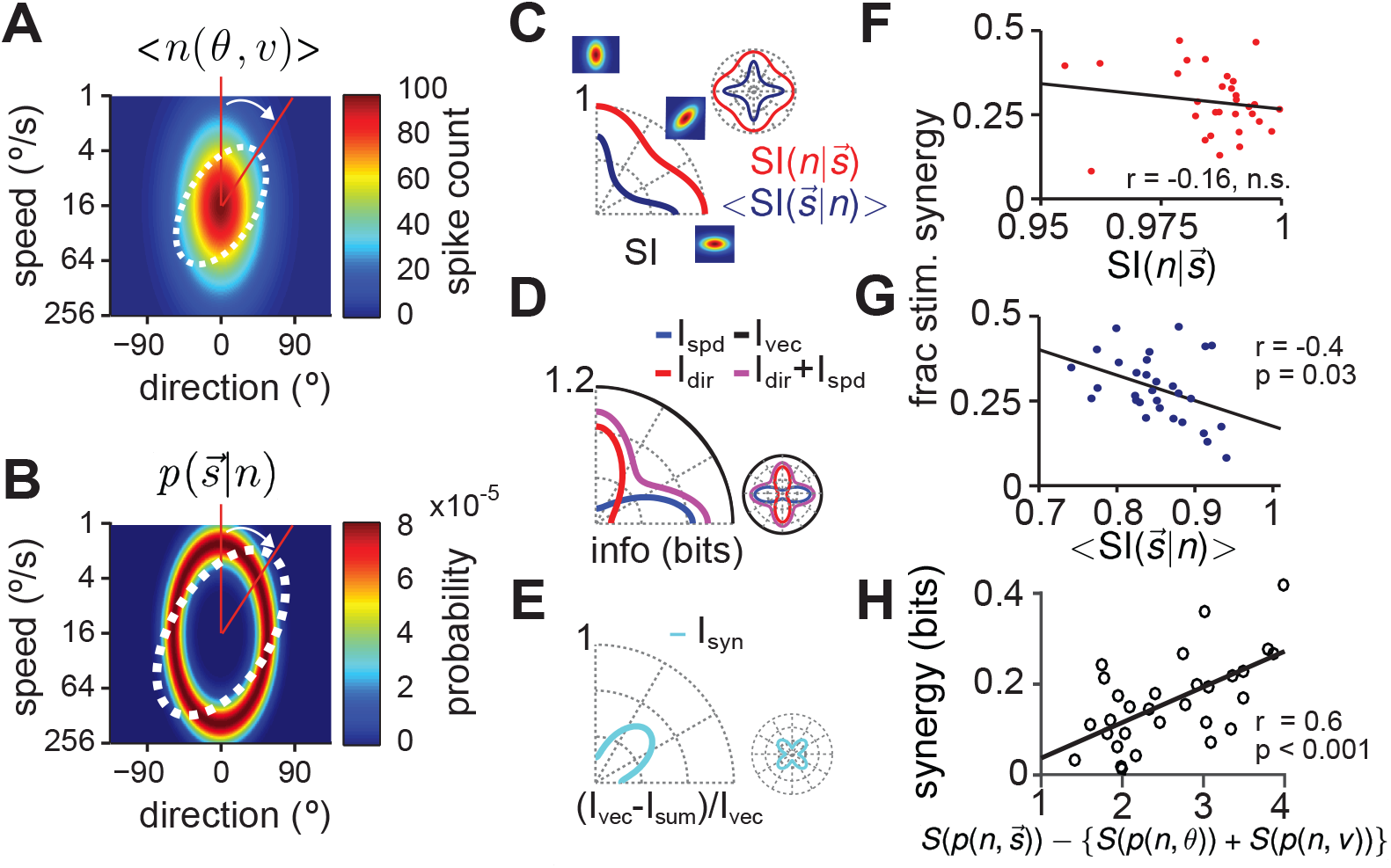
Contributions of conditional response and stimulus distributions to stimulus synergy. **A.** An idealized 2D direction-speed tuning function, 〈*n*(*θ, v*)〉. When the long axis of the function is oriented along cardinal directions, direction and speed tuning functions are separable. At intermediate angles, the response to each motion component is not independent of the other. **B**. As the tuning function is rotated, the conditional stimulus distributions, 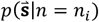 rotate with it, changing their separability. **C**. The separability index (SI, see text) is modulated as a function of rotation angle. The SI of 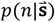 (red) is consistently higher than the SI of 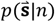 (blue). **D.** Mutual information between Poisson spike counts, generated with the tuning function in A, and stimuli. Information about direction (red) and speed (blue) are modulated by tuning bandwidth, which changes as a function of rotation angle. The information about the 2D motion vector (black) is constant under rotation, and is always larger than the sum of the information about each stimulus component (magenta). Radial axis has been normalized such that the vector information is unity. **E**. Fractional synergy as a function of rotation angle. Stimulus synergy is maximal when the separability of both 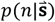 and 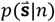 are at their minima, but stimulus synergy is non-zero around the circle. **F.** SI of 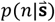 is not predictive of fractional synergy. **G.** <SI>, defined as 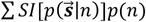, is correlated with fractional synergy in model units generated from recorded MT neurons. **H.** Synergy is correlated with the difference between the entropy of 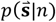 and the entropy of *p*(*θ|n*) *p*(*v|n*).

### Stimulus synergy is modulated by tuning function separability

Not all sensory neurons will have separable tuning functions, in which case stimulus synergy might arise from the form of the likelihood, 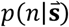, as well as the posterior, 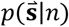. In MT, units have a distribution of seperabilities that impact synergy in population estimates. To consider more generally how tuning function inseparability could contribute to stimulus synergy, we rotated separable Gaussian direction and speed tuning functions (SI=1) matched to our MT data and calculated synergy at each angle (Figure S2). Rotating 2D tuning functions in the direction-speed coordinate frame alters the degree of separability between direction and speed tuning (blue trace, Figure S2C), and likewise the separability of 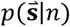 (red trace, Figure S2C, 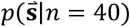 shown) which rotates along with 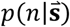 and whose SI depends on the count Fig S1F). SI values are lowest at oblique orientations, and the SI of the posterior is always lower than the SI of the tuning function. Information about direction (red trace, Figure S2D) and speed (blue trace) and their sum (magenta trace) also rises and falls with rotation angle because direction and speed tuning bandwidths vary, however information about the 2D motion vector (black trace, Figure S2D) remains constant. Synergy values are highest at oblique orientations (cyan trace, Figure S2E) when tuning is least separable, however, the degree of synergy is not well predicted by separability alone. The correlation between tuning SI and the level of synergy is not statistically significant (Figure S2F). While the separability of the mean posterior 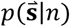 is anti-correlated with the degree of stimulus synergy (r = −0.4, p = 0.03, blue symbols Figure S2G), the difference between the likelihood and posterior entropies is the best predictor (Figure S2H). More information can be encoded about a multidimensional stimulus if the entropy of 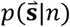 is lower than the entropy of *p*(*θ|n*) *p*(*v|n*). Indeed, the entropy difference is highly correlated the degree of stimulus synergy we observe in each MT neuron (Figure S2H).

### Stimulus synergy can also arise from monotonic tuning functions

**Figure S3.**
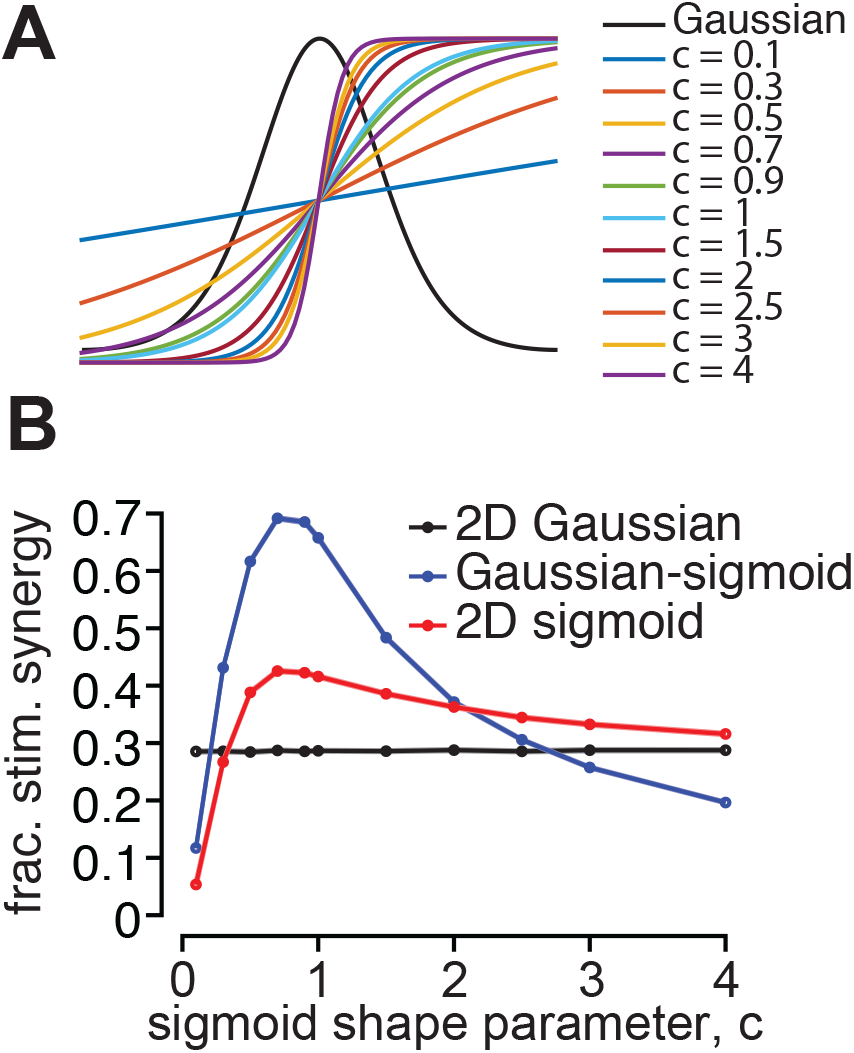
Stimulus synergy is enhanced by monotonic tuning functions. **A.** We compared stimulus synergy arising from Gaussian tuning, monotonic (sigmoidal) tuning, and a combination. Sigmoidal functions varied in steepness with the value of a shape parameter, c. **B**. Fractional stimulus synergy as a function of the sigmoid shape. The black trace represents synergy with 2D Gaussian tuning, blue trace Gaussian tuning for one feature and sigmoidal tuning for the other, and the red trace 2D sigmoidal tuning. Stimulus synergy is enhanced by sigmoidal tuning for all but the steepest and shallowest sigmoids.

To this point our analysis has focused on neurons with Gaussian-like tuning functions, but bell-shaped tuning is not necessary for stimulus synergy. For example, visual neurons that have Gaussian tuning for direction or orientation might have monotonic tuning for contrast or binocular disparity. We computed the degree of stimulus synergy when one parameter has Gaussian tuning, and another sigmoidal (monotonic) tuning, and when both have sigmoidal tuning. We find that stimulus synergy is enhanced by sigmoidal tuning across a wide range of steepness (Figure S3). For all but the shallowest, nearly linear tuning functions (blue trace, Figure S3A), and the steepest transitions from low to maximal firing rate (yellow, purple traces, Figure S3A), monotonic tuning enhances stimulus synergy with respect to 2D Gaussian tuning (black line, Figure S3B). None of the monotonic tuning functions eliminate synergy altogether. This is also true when tuning for both stimulus features is monotonic (red trace, Figure S3B).

### Stimulus correlations and stimulus synergy

We have demonstrated that stimulus synergy can occur in the absence of correlations between stimulus features because our stimulus presentation ensured equal likelihoods. Under these conditions, interactions between direction and speed estimates observed in MT and pursuit behavior had to arise from internal computation. In the natural world, however, the stimulus features of interest may not be independent, i.e. *p*(*s*_1_, *s*_2_) ≠ *p*(*s*_1_)*p*(*s*_2_), which could impact the level of observed stimulus synergy. Correlations between stimulus features do not change the definition of stimulus synergy, but they do change the joint distribution between responses and inputs, and thus affect mutual information and synergy. If the stimulus is 2D with 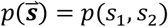, stimulus synergy is defined as 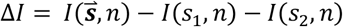 and is equal to the negative of the co-information between three random variables, −*I*(*n, s*_1_, *s*_2_), also termed the interaction information. Note that the definition of a coding synergy is −*I*(*s*_1_, *n*_1_, *n*_2_), where *n*_1_ and *n*_2_ are features of the neural response like unit identity or spike times^10^. Unlike the mutual information between two variables, interaction information can be positive or negative, resulting in redundancy or synergy. Using information theoretic identities, one can show that the definition of stimulus synergy is equivalent to Δ*I* = *I*(*s*_1_, *s*_2_|*n*) – *I*(*s*_1_, *s*_2_), the difference between the mutual information between two stimulus features having observed a neural response and their mutual information in the environment. Synergy is present whenever observation of neural activity increases the magnitude of correlations between estimates of the stimulus. If *s*_1_ and *s*_2_ are uncorrelated, their mutual information is zero, and stimulus synergy is maximized. Correlations between *s*_1_ and *s*_2_ imply that *I*(*s*_1_, *s*_2_) > 0, reducing stimulus synergy. In practice, high levels of correlation are necessary to eliminate the synergistic effect of multidimensional tuning. Figure S4 shows how *I*(*s*_1_, *s*_2_), *I*(*s*_1_, *s*_2_|*n*), and stimulus synergy vary as a function of linear correlations between two Gaussian distributed random variables for a neuron with MT-like tuning. The mutual information between *s*_1_ and *s*_2_ increases gradually with increasing correlations (positive or negative) while the conditional information remains constant, dominated by the tuning functions. Stimulus synergy (Δ*I*) is present for correlations less than |*ρ*| < 0.7. The level of motion correlations in natural moving scenes, for example, is substantially lower, |*ρ*| < 0.15^35^, a regime where correlations have a negligible impact on stimulus synergy.

**Figure S4.**
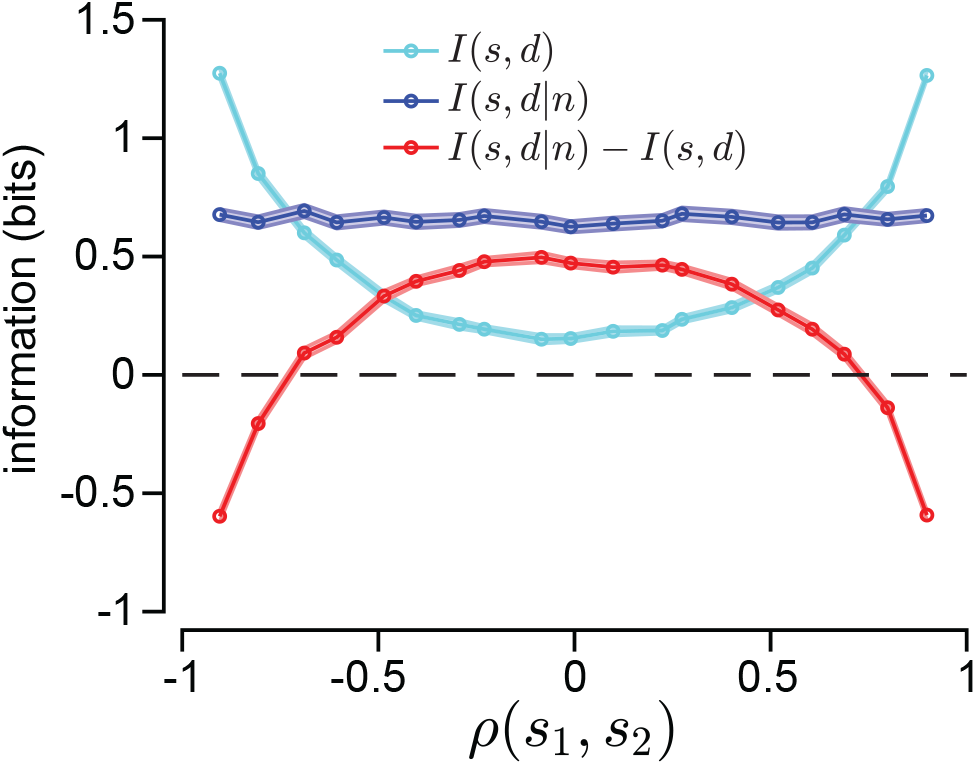
Impact on stimulus correlations on synergy. We generated Gaussian distributed directions and speeds with linear correlations *ρ*(*s*_1_, *s*_2_)that ranged from −1 to +1. Using the mean 2D MT tuning function, we generated Poisson distributed count samples and then computed the mutual information conditioned on particular count values. The expectation value of the conditional information (*I*(*s*_1_, *s*_2_|*n*) (dark blue trace) remains constant across the range of input correlation levels. *I*(*s*_1_, *s*_2_) (cyan trace) increases with the magnitude of correlation. Information doesn’t go to zero with *ρ* = 0 due to small residual correlations in stimulus samples. Stimulus synergy is the difference between these quantities, Δ*I* = (*I*(*s*_1_, *s*_2_|*n*) – *I*(*s*_1_, *s*_2_) (red trace). Positive values, i.e. synergy, persists for *ρ* < 0.7 with higher correlation levels creating redundancy.

Input correlations will have little impact at the population level and in behavioral output. Behavioral, or output, synergy can be written 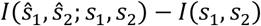 and population synergy by 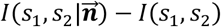. Input correlations will shift both terms by a constant that is small compared to the first terms. Note that the difference between population synergy and behavioral synergy will not be affected – the *I*(*s*_1_, *s*_2_) terms will cancel, leaving the behavioral constraints on decoder efficiency unaltered by correlated inputs. Writing stimulus synergy this way emphasizes that it is a means of quantifying both neural sensitivity to a conjunction of stimulus features and the potential downstream consequence of ignoring the correlations induced by multidimensional tuning for decoders of neural activity and thus for behavior.

## Discussion

Multidimensional tuning creates a credit assignment problem − along which dimension did a stimulus change to create the observed modulation in activity? The fact that multidimensional maps are the rule rather than the exception throughout the brain might arise from anatomical wiring or metabolic efficiency constraints, hierarchical processing efficiency, or behavioral convenience, yet we show that multidimensional tuning also affords a computational advantage in encoding information even in the absence of inseparable tuning functions. The fact that stimulus synergy is observed in pursuit behavior has several implications for how sensory activity is read-out to guide behavior. First, it implies that sensory stimuli are decoded in the dimensionality of their representation, i.e. as a motion vector rather than independent direction and speed estimates from MT. Second, it implies that the decoders are efficient, maximizing information output. These results add to a growing body of work demonstrating efficient information transmission from visual motion estimates to pursuit eye movements^19–21,36^.

Unlike more familiar forms of coding synergy, stimulus synergy does not depend on subtle aspects of feature selectivity or spike timing, but rather emerges from multidimensional tuning itself. Therefore, we expect that stimulus synergy will impact information representation and recovery in any area of the brain with tuned (e.g. Gaussian, ramped, sigmoidal) input-output functions. Like MT, neurons in primary visual cortex (V1) are tuned for a small number of visual features. Grunewald and Skoumbourdis reported that V1 neurons tuned for binocular disparity and orientation encoded more information about the features jointly than singly, although their focus was the impact of tuning diversity within V1 on coding^37^. Recent studies of prefrontal cortical neurons credit their high-dimensional representations with enhanced cognitive performance on decision-related tasks^38,39^. Our work provides a theoretical framework for understanding how multidimensional representations, whether in sensory or higher association areas, could benefit the many behaviors that depend on multidimensional estimates from cortical populations.

### Origins of stimulus synergy

Coding synergies can arise from neural diversity within nominally redundant coding pools^11,13^, or from temporal patterning in spike trains^10,40^ such that the pattern encodes more information than the summed response. MT neurons do demonstrate classic coding synergies, largely arising from diversity in rate dynamics^13^ and some from temporal patterning in spiking^10^, but neither is the origin of stimulus synergy. Even when the tuning for each stimulus feature is separable, multidimensional tuning creates a divergence between the conditional response distribution for an n-dimensional stimulus vs. its component parts, and it is this nonlinearity that creates stimulus synergy. Of course, neurons that response only to a particular conjunction of stimulus features like the canonical face neuron also display substantial stimulus synergy.

### Implications for decoding multidimensional stimuli

A potential downside to multiplexing stimulus features is that unidimensional components cannot be decoded efficiently using only knowledge of the marginal statistics of the stimulus conditioned neural response. In this setting, efficient decoding requires either joint decoding of stimulus variables (decoding the stimulus in the dimensionality of its representation) or an additional circuit mechanism that applies an appropriately tuned (i.e. stimulus dependent) non-linear operation capable of converting the multiplexed representation into a representation that is devoid of stimulus synergy, an operation known as marginalization in the probabilistic reasoning literature. The latter possibility is not consistent with our behavioral data that displays substantial stimulus synergy. Marginalization is a necessary component of inference in general and for smooth pursuit in naturalistic settings in particular. This is because targets have visual features such as shape and color that affect activity in the visual system but are not the quantities of interest (i.e. direction, speed) when generating tracking eye movements. Behaviors such as pursuit demonstrate the brain has likely solved this problem. The critical cortical computation for both decoding and marginalization is a tuned form of divisive normalization, a circuit level operation that is ubiquitously observed in cortex^41,42^. Thus, while a multiplexed neural code can present difficulties for downstream decoders, they are not the kinds of difficulties that are uncommon to cortical computation.

From an empirical perspective, the presence of stimulus synergy in sensory cortex is highly advantageous since synergy that arises from correlated errors is also present in behavior. Indeed, high degrees of synergy in sensory cortex coupled with the assumption of efficient inference and decoding leads to the behavioral prediction that, not only should errors be strongly correlated, but that the structure of those correlations should mimic the correlations of the response conditioned posterior that drives coding synergy. This places strong constraints on the set of candidate neural decoders that the brain uses to generate behavior, and it creates a rich substrate for future investigation.

### Are sensory responses tuned to maximize multidimensional stimulus information and stimulus synergy?

If information efficient representation is a primary objective in sensory map design, are cortical sensory neurons encoding the ideal number of stimulus features? That is a difficult question to answer because it is unclear whether we have fully described feature selectivity in any cortical area, and the predicted level of stimulus synergy will depend critically on that tuning among other factors. Most MT neurons display strong, tuning for direction and speed, but many are also modulated by other visual features, such as spatial frequency^16^, acceleration (Lisberger and Movshon 1999), or binocular disparity^44^ -- although it is unclear the extent to which MT responses are tuned to these other features. For the average tuning bandwidth, we measured in our MT sample, 2-3 dimensions correspond to a peak in total stimulus information and to stimulus synergy. While this result is suggestive, a comparative study of stimulus synergy in neurons with a large variety of feature selectivities and dimensionalities will be needed to draw any conclusion about its generality. If maximizing stimulus information about each stimulus feature were the brain’s sole objective, we should observe a preponderance of highly redundant and highly selective single feature sensory neurons. To the contrary, many if not most sensory maps represent more than one stimulus dimension. The reason is likely that neurons and the spikes they generate are metabolically expensive, and cranial space is at a premium, such that energy and wiring savings may outweigh the modest information loss. It is also possible that multidimensional tuning could have computational benefits such as simplifying hierarchical processing. Certainly at the population level, the tuning bandwidths that optimize encoding depend on other factors such as response overlap, heterogeneity, pairwise correlations, and the coding symbol of interest^1,2^.

It is also interesting to note that, in our simulations, synergy and efficiency are coupled as population size increases. Specifically, in the presence of limited input information, synergy is highest when the mutual information between stimulus and response begins to saturate to its maximal value. This is precisely the point at which adding more neurons to the population begins to have less and less of an effect on errors. While more work on the link between synergy and efficiency in this setting is required to draw strong conclusions regarding their relationship, this result suggests that high degrees of stimulus synergy in behavior may be a hallmark of a neural code that is optimized to maximize information transmission (or minimize errors) using the fewest spikes or neurons.

Our work demonstrates that stimulus multiplexing in sensory areas can confer a computational advantage, and it shows that behavior can benefit. Stimulus synergy should influence how we think about sensory representations and their readout, and it highlights what fundamental coding features neuroscientists can miss when isolating single stimulus features in experimental design.

## Supplementary Material

## Appendix 1. MT 2D tuning function separability

MT firing rates are strongly modulated by both the direction and speed of motion. MT direction-speed tuning functions are ovoid in shape (with standard stimulus units) and oriented such that direction tuning does not change as a function of stimulus speed, and vice versa (Figure S1C; c.f. Figure 1E). We used singular value decomposition to create a separability index (SI) (see Methods) to quantify the degree of nonlinear interaction between direction and speed tuning. SI values for the 6 example neurons are indicated in the upper right corner of each panel in Figure S1C, and population data are shown in Figure S1E (red bars). Perfect separability (SI=1) indicates that the average spike count for any direction-speed combination can be predicted from the 1D tuning functions alone, i.e. 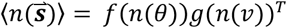. MT SI values cluster near complete separability (SI=1), with a single mode describing 98±1% (mean ± SD, n=30 units) of the variance in the shape of the conditional response distributions. Second SVD mode values were below statistical significance for all units. While the tuning functions did not display nonlinear direction-speed interactions conducive to coding synergies, the conditional stimulus distributions, 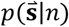, were highly non-separable. We plot 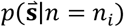 for the same 6 MT neurons in Figure S1D, where *n_i_* is the mean spike count observed for a stimulus at 16°/s and 60° from the neuron’s preferred direction. Their shapes make it apparent that the marginal distributions of directions *p*(*θ*|*n*) and speeds *p*(*v*|*n*) that give rise to a count *n* cannot be multiplied to yield 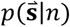, i.e. 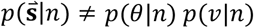. The sample mean SI value for 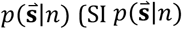, blue bars, Figure S1E), is lower (0.87±0.09 SD, n=30 units) than for 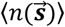 (SI 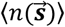, red bars, 0.99±0.007 n=30 units, p<10^-5^, Kruskal-Wallis one-way ANOVA). The inseparable form of the posterior distribution provides a substrate for stimulus synergy that is not apparent in the tuning function.

**Figure S1.**
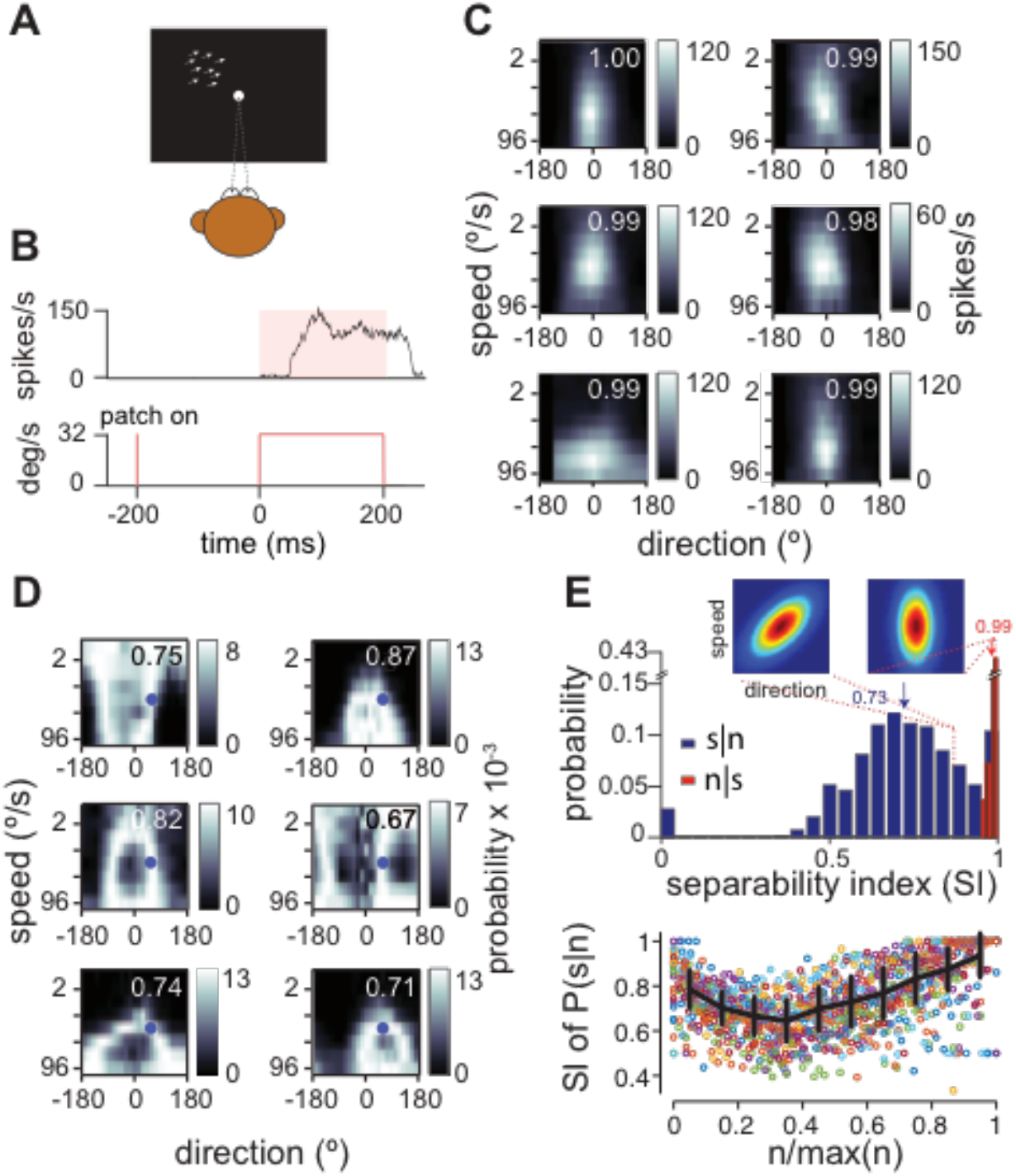
Separable direction-speed tuning functions, inseparable posterior distributions in MT. **A**. Monkeys fixated while random dot kinematograms moved coherently in each unit’s receptive field. **B**. (bottom) Dots appeared and remained stationary for 200 ms, then moved in one of 12 or 16 directions and 8 speeds for 200 ms, then were stationary for another 300 ms. (top) Example peri-stimulus-time-histogram (PSTH) for one motion condition sampled at 1000 Hz and summed over sliding 10 ms bins. **C**. 2D directionspeed tuning functions for 6 MT units. Panels show the conditional mean spike count 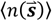 from 0-200 ms after motion onset. Separability indices (SI, see text) listed in upper right corner. **D**. Response conditioned stimulus distributions, 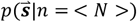, where <*N*> is the mean spike count observed for a stimulus at 16°/s and 60° from the neuron’s preferred direction, for the same 6 units. SI values are indicated as in C. **E.** Histogram of separability indices across MT sample for both the conditional mean count (SI 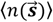, red bars) and the conditional stimulus distribution (SI 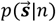, blue bars). A perfectly separable tuning function (right inset) has an SI index of 1. Rotating the tuning function by 45° (left inset) creates a maximally inseparable 2D tuning function with the same 1D tuning functions (SI = 0.88). **F**. SI of 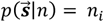 for all *n_i_* from 1 to *n_max_*. Each dot represents a single count value for a single unit. Black trace represents the population mean, SD. Posterior separability is lowest for intermediate count values when the most likely combinations of direction and speed lie on a ring. SI values are high when the count is maximal when only a single direction, speed combination is most likely forming a point. SI values for the lowest counts are also higher because the posterior distribution is more diffuse than ring-like.

The inseparability of the posterior stimulus distribution depends on the spike count (Figure S1F). If the count is maximal, near the peak of the 2D tuning curve, then the posterior is approximately a point in the stimulus space, neglecting noise. A point necessarily has an SI of 1. However, for all other response values, SI values are lower such that the average SI is well below 1.

## Data Availability Statement

Data will be made available upon reasonable request.

## Code Availability Statement

All software used for data analysis is available upon reasonable request.

## Acknowledgements

We thank S.E. Palmer, D.J. Schwab, and N Brunel for early discussions about the analysis. We thank T. Mukherjee and Bonnie Bowell for assistance with animal care and data collection, and the veterinary staff at Duke University and the University of Chicago. M.V.M. was supported by the Luckhardt Endowment Fund at the University of Chicago. Research was supported by grants to L.C.O. from the Alfred P Sloan Foundation, Whitehall Foundation, Brain Research Foundation, NIH NEI EY023371, and NSF IOS 145704.

## Methods

### Experimental Methods

We recorded single-unit MT responses and eye movements in 3 adult male rhesus macaques (*Macaca mulatta*). Prior to these experiments, the monkeys had been extensively trained in pursuit and fixation tasks. We fit each monkey with a head restraint and sclera-implanted eye coil under general anesthesia, using sterile surgical technique and postoperative analgesics. All study methods were approved in advance by Duke University’s and the University of Chicago’s *Institutional Committee for the Care and Use of Animals* (IACUC) and were in full compliance with guidelines from the National Institutes of Health.

### Visual stimuli

For both the physiology and pursuit experiments, monkeys viewed bright visual stimuli against the dark screen of a fast CRT display (SGI GDM-FW9011 or Dell X1250) at a 100Hz frame rate and 1024×768 pixel resolution at a distance of 48.5 or 51.5 cm in a dimly lit room. Full field stimuli subtended 53° by 36° (SGI) or 36° by 33° (Dell). For physiology experiments (Figure S1A), stimuli consisted of bright random dot kinetograms (1 dot = 2 pixels; 2 dots/deg^2^) with constant speed and direction displayed against the dark background of the screen. Dots moved coherently in a stationary rectangular aperture matched to the preferred size of the recorded cell. Pursuit experiments employed the same dot motion stimuli with matching center of mass motion such that the dot pattern translated across the screen as a cohesive object. For some pursuit experiments we used a uniformly illuminated circular spot target subtending 0.4° of spatial arc (Figure 3A). Trial time courses are defined for the physiology and pursuit experiments below.

### Physiological recording

We recorded single units in area MT using a 3-electrode array (TREC, Germany) of high-impedance microelectrodes. Our data acquisition system (Plexon Omniplex, Blackrock Cerebus) sampled neural activity at 30kHz, and stored each channel’s waveform for offline spike sorting. We identified area MT based on stereotactic coordinates along with online testing of direction and speed tuning and classical receptive field size. We confirmed our isolation of single units through principal component analysis of spike waveforms in tandem with inspection of inter-spike interval (ISI) distributions. We initially mapped each unit’s motion selectivity online using 400 ms steps of full field (50° by 50°) coherent dot motion in 8 directions (45° increments) and 8 speeds (1-96°/s in powers of 2). Using the coarsely determined preferred speed and direction, we mapped the receptive field (RF) location with 400 ms steps of dot motion in 5°-square apertures that appeared at pseudo-random positions at 5° spacing on an 8 by 6 grid, covering 20° by 15° of visual space. We interpolated the RF center and remapped the RF in a similar way, using 2°-square apertures at 2° spacing on an 8 by 6 grid, covering 8° by 6° of visual space. We tested a range of aperture sizes (1-40°) and selected the size that drove the maximal firing rate to use for our experiments. Finally, we remapped the unit’s direction tuning with 400 ms of dot motion at 15° intervals before proceeding with the experiment.

We organized the physiology experiments into trials of 1-2s duration during which monkeys maintained fixation. After a 200 ms fixation interval, we projected random dot patterns in square apertures scaled to the preferred size in each unit’s RF. The dots remained stationary for 200 ms to allow firing rates to return to baseline, then stepped to move in a constant direction and speed for 200 ms, then remained stationary for another 200 ms (Figure S1B). Inter-trial intervals were approximately 2-3s. We aligned the stimuli set with each unit’s preferred direction (see above) and recorded responses to motion at 45° increments with around the circle at speeds of 1, 2, 4, 8, 16, 32, 64, or 96°/s, comprising 64 motion stimuli. Directions of ±75°, ±60° were added for 4 cells to verify that the distribution of chosen directions did not affect analysis (96 stimuli). For 16 units, trials were concatenated to include 3 motion steps in different directions at a single speed, each step interleaved with a 200 ms stationary period. Other neurons in the data sample were recorded with a single stimulus motion per trial as described above. Trial types were pseudo-randomly interleaved in blocks. Monkeys were rewarded following each trial for maintaining fixation within 2° of the fixation spot. We presented each stimulus an average of 36 times (range 15-87 repetitions). Single unit isolation was confirmed with Plexon’s off-line spike sorter using principal component analyses of the recorded waveforms. We only report data from clearly isolated waveforms. 30 of 40 recorded units met our criteria for isolation.

### Pursuit experiments

We trained 3 adult male macaques (M1-3) to fixate and pursue moving targets for a juice reward prior to data collection. We sampled the horizontal and vertical position of one eye at 1 ms intervals and then filtered, differentiated, and digitized the signals. During pursuit experiments, animals were required to maintain fixation within 2° for 500-1000 ms to begin a trial and during the final 200 ms of the pursuit interval and for 400 ms at the end of the trial. Accuracy windows were relaxed during pursuit initiation. Pursuit trials were ~2 s in duration. The fixation spot appeared at the center of the screen for a randomly selected interval from 500-1000 ms. The fixation spot then disappeared, and a target appeared ~3° eccentric to screen center and immediately moved at a constant direction and speed toward the point of fixation. For each monkey, we selected the eccentricity of target appearance to minimize the occurrence of saccades during pursuit initiation. Targets moved for 600 ms then jumped 1° in the direction of motion, remaining stationary a final fixation period of 300 ms (Figure 5A, bottom). Spot targets moved in 14 directions (0°, ±10°, ±20°, and ±30° relative to leftward and rightward motion) and 5 speeds (10, 15, 20, 25, or 30°/s) for 70 different target motions (Figure 3B). We did not analyze trials with saccades or eye blinks during a 500 ms interval from target motion onset. We collected 33-128 repetitions (mean 94) of each target motion for each pursuit data set across multiple sessions.

To measure the period over which pursuit is under open-loop control, we compared eye velocities during normal pursuit to trials in which we forced the control loop to remain open by moving the target along with the eye (Lisberger and Westbrook 1985; Osborne et al. 2007). By adding the eye velocity to the target velocity (20°/s), we maintain a constant target speed on the retina. The eye velocities for the stabilized trails will diverge from normal, un-stabilized pursuit at the time point at which extra-retinal feedback signals produce a measurable impact on eye speed. We defined the duration of the open loop interval as the time point with respect to pursuit onset that mean eye speeds diverged by 1 SD. The open-loop interval was 146 ms (M1), 135 ms (M2), and 138 ms (M3), consistent with past studies^19,21,45^.

## Quantification and Statistical Analysis

### Separability analysis in MT units

We used singular-value decomposition (SVD) to characterize the degree of separability of the 2D direction-speed tuning functions of our MT sample. The fractional power of the first singular value *ui* with respect to the other *i* modes, i.e. 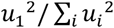 quantifies the degree to which the 2D tuning function, *f*(*θ, ν*) is captured by the outer product of a direction-only and speed-only tuning curves, *f*(*θ*) *g*(*ν*)^*T*^. This value defines a separability index (SI) that ranges from small values to 1^29,37^. An SI of 1 indicates that variance in the filter form is perfectly captured by a single mode and hence, its spacetime separability is exact. Randomly scrambling MT 2D tuning functions, which destroys tuning, produced SI values of ~0.4 whereas rotating the 2D tuning function to an oblique angle lowers SI to ~0.9.

To investigate the role of tuning function separability to stimulus synergy (Figure S1) we created data-driven model MT neurons with homogeneous Poisson spike statistics so we could manipulate separability. We fit 1D Gaussian functions to the peak firing rate, tuning width and baseline firing rate each neuron’s spike count in a 200 ms window with respect to stimulus motion onset. We used responses to preferred speed to fit the direction tuning curve and vice versa. We measured speed tuning in log_2_ units of stimulus speed. The outer product of each model neuron’s direction and speed tuning curves provided the mean spike rate λ for a Poisson count distribution for each stimulus condition 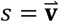:

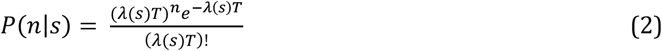

We interpolated the 2D tuning curve to 5° spacing around the circle and 0.2-unit spacing in log_2_ speed (i.e., 5, 5.2, 5.4 in log_2_ spacing = 32, 36.8, 42.2 in degrees/s). We manipulated the direction-speed separability by rotating the ellipsoidal 2D tuning curve about its preferred stimulus in 1° steps. At a rotation angle of 0°, 90°, 180° or 270° the tuning curve is perfectly separable, with an SI of 1. At a rotation angle of 45°, the tuning curve has an SI of 0.88, and the direction and speed tuning functions are not independent, i.e. inseparable^21^. We use the definition that a separable distribution is one where the second SVD mode does not reach statistical significance (p < 0.01) using a permutation test. We randomly permuted the elements of each tested matrix 1000 times and used the distribution of the separability values to determine whether the second SVD mode was above chance. Rotating a separable 2D Gaussian tuning function by >20° increases the second SVD mode above chance levels. The SI at a 20° rotation is 0.95, so we classify any SI below 0.95 as inseparable.

### Information measures

We quantified the relationship between motion stimuli and MT response with the mutual information^19,21^, which captures in bits the average reduction in uncertainty about stimulus identity from the observation of a neuron’s response. We defined the neural response as the total number of spikes fired in a time window beginning at motion onset up to some time, *T* – i.e. the cumulative spike count. The rate at which the count increases changes over time based on dynamics in the firing rate and is highly dependent on the direction and speed of the motion stimulus. We sampled the cumulative count at 1 ms resolution to form response distributions and entropies. We computed information values (Eqn. 4) with 50 random draws of different fractions of our data sample, computing the means and standard deviations (SD) of all quantities. We corrected for finite sample effects by inspecting and fitting a linear function to the information values as a function of 1/(no. of trials) in each fraction of the sample and extrapolating to an infinite sample size^26,27^. All reported values had finite size corrections of less than 10%.

We used the same method to compute the mutual information between the eye velocity vectors and target motion vectors over time. We sampled the horizontal and vertical components of the eye velocity at 1 ms intervals and analyzed the movements starting from pursuit onset for each target condition, estimated from trial-averaged responses. At each time step we discretized the range of observed eye directions and speeds across all target conditions into bins with near-equal occupancy (adaptive binning). Behavioral information calculations then proceeded identically to the methods described for neural data. Because of the limited stimulus entropy, the direct method results in lower bit levels than translating eye velocity SNR to information^20^.

### Stimulus synergy in single neurons

Synergy is typically defined as the excess of information available from a pattern of responses (across cells or over time) compared to the sum of the information contributed by each^10^. Here we define stimulus synergy to be an excess of information encoded about a multidimensional stimulus vector compared to its components. Note that the definition of stimulus synergy does not require that the stimulus be a vector in the mathematical sense - stimulus features are defined by the experimenter. For a stimulus vector with *N* elements, the information difference is given by:

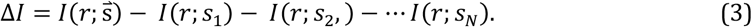

When Δ*I* > 0 coding is synergistic. If Δ*I* < 0 coding is redundant, and if Δ*I* = 0 coding is independent. For each si, the mutual information is defined as

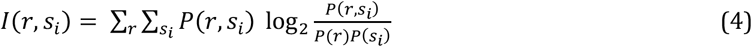

If 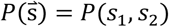, then

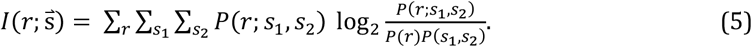

Substituting Equations 4 and 5 into 3 and consolidating terms yields

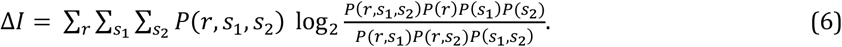

If Δ*I* > 0, then Eqn. 6 defines stimulus synergy for a 2D stimulus. The RHS of Eqn. 6 is equivalent to −*I*(*n, s*_1_, *s*_2_).

We can isolate contributions from coding non-linearities and stimulus correlations to Δ*I* by exploiting the symmetry of mutual information:

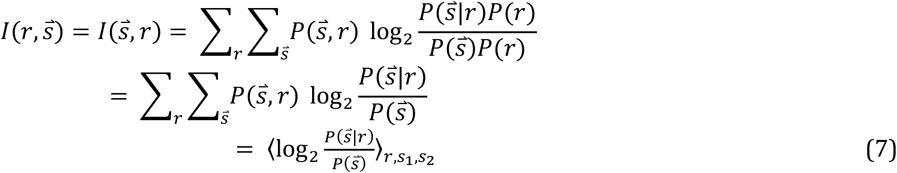

where the <…> indicate an expectation value. Repeating this operation on the marginal distributions, Equation 3 (or 6) becomes

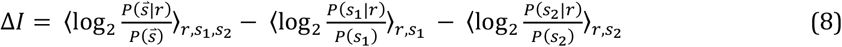

Rearranging terms algebraically gives

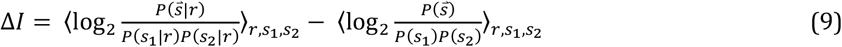

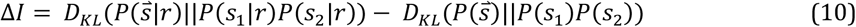

where the two *D_KL_* terms in Equation 10 represent Kulback-Leibler divergences. The first term expresses the extent of decoding nonlinearity, and the second term grows with correlations between stimulus features. Like more traditional forms of synergy, stimulus synergy arises from a divergence between a vector distribution and its scalar components, but that vector represents features of the stimulus rather than of the neural response. Each term in Equation 10 is non-negative, so stimulus synergy is zero only when 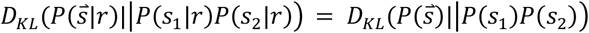. This can occur trivially when *s*_1_ and *s*_2_ are independent both prior to observation of neural activity and after observation of neural activity. Alternatively, interpreting KL divergence as a generalized correlation coefficient, we could say that synergy is zero when observation of neural activity does not change the structure of the correlations between *s*_1_ and *s*_2_.

In order to isolate the impact of non-linearities in stimulus decoding (the first term in Equation 10), our experiments presented stimulus features in an uncorrelated manner such that the second term vanished. Our stimulus was a 2D vector, 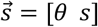, which specifies direction *θ* and speed s presented such that 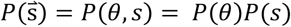. A priori correlations between stimulus features will impact both terms in Eqn. 10, and could potentially create negative values of ΔI (i.e. redundancy).

#### Stimulus synergy in population responses

At the population level, the response becomes a pattern of spike counts across neurons that is typically too highdimensional to sample experimentally, so we used data-driven simulations to model MT population responses. For each model unit, we generated spike counts with homogeneous Poisson statistics from a separable rate tuning function:

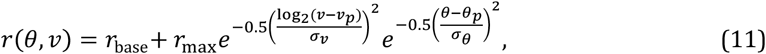

where *r_max_, v_p_, σ_v_*, and *σ_θ_* were set to be roughly comparable to our sample of neurons and to published ranges. Each unit had a randomly selected tuning parameters, and background and peak firing rates. Base firing rates (*r*_base_, mean = 6 spikes/s, SD = 1 spikes/s) were Gaussian distributed and rectified to be >0. Peak firing rates (*r_max_*, mean = 100 spikes/s, SD = 0.5*mean) were Gamma distributed, direction tuning width (*σ_θ_*, mean = 40°), SD = 0.3*mean, offset = 5°), and speed tuning width (*σ_v_*, mean = log_2_(16°/s), SD = 0.3*mean, offset = 0.5) were also Gamma distributed. The distribution of preferred directions had an overrepresentation near cardinal directions after Xu et al., 2006^46^. The distribution of preferred speeds was taken from Priebe et al., 2006^47^. We approximated the narrow distribution of MT separability indices by introducing small, randomly generated correlations between direction and speed tuning functions, equivalent to very small rotations of the 2D tuning function. The mean separability index was >0.99. Population sizes ranged from 2^4^ to 2^11^ units.

We generated noise correlations between units based tuning similarity. For a pair of neurons with preferred directions (*θ_i_, θ_j_*), direction tuning bandwidths (*σ_θi_, σ_θj_*), preferred speeds (*v_i_, v_j_*), and speed tuning bandwidths (*σ_vi_, σ_vj_*), the correlation matrix element is given by:

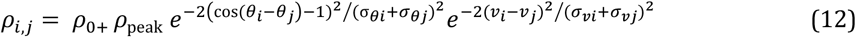

Figures were generated with *ρ*_0_ = 0.05 and *ρ*_peak_ = 0.6 with a population average value of *ρ_i,j_* of ~0.15. To ensure that the correlation matrix was positive definite, we replaced all negative eigenvalues of the matrix generated from known pairwise correlations with a small positive number (epsilon = 0.001). In the limit as epsilon goes to zero, this procedure is equivalent to finding positive definite semi-definite that is nearest (with respect to the Frobenius norm) to the original pairwise correlation matrix. We then normalized the diagonal and scaled the correlation matrix by the stimulus dependent mean firing rate to create a response covariance matrix, **Σ**. We perturbed **Σ** slightly by adding information-limiting ‘correlations’^32^. We used the precision of pursuit and perception to set the level of information-limiting correlations. On this time scale, motion discrimination thresholds are ~3° and ~10% of the average speed^21^. We took these quantities to be the inverse Fisher information for direction and speed (described below) and scaled the level of information limiting correlations (*ρ*_IL_) in units of these values. Figures 4 and 5 were generated with *ρ*_IL_ =2. These adjustments perturbed the original correlation values by less than 1%.

The implication of stimulus synergy is that a joint decoder that estimates the vector of stimulus features can recover more information than a set of marginal decoders that each estimates the value of a single stimulus feature. We tested that hypothesis at the population level by comparing the linear Fisher information from the joint stimulus distribution compared to its marginals. In our case the Fisher information is a 2×2 matrix:

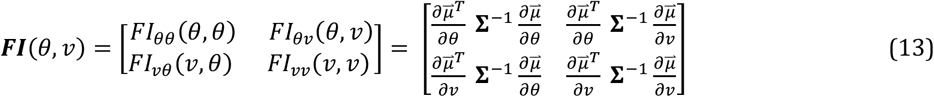

where 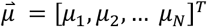 is the population vector of average spike counts in a time window for an *N*-cell population for a specific stimulus (*θ, v*), **Σ** is the *N×N* covariance matrix of count fluctuations for that stimulus (i.e. 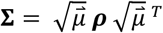), and the superscript T indicates a transpose operation. Equation 13 drops the stimulus specification (*θ, ν*) for each term for simplicity. FI values for the marginal stimulus distributions are scalar quantities computed in a similar manner, but with respect to a single stimulus feature.

Analogously to mutual information in Equation 3, stimulus synergy is present for a population response at a stimulus (*θ, v*) when

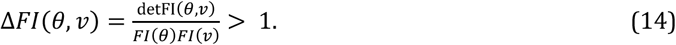

### Decoding models

After constructing the population response, we employed several decoding models to estimate stimulus direction and speed both separately and jointly. Fisher information was then computed between the stimulus and the estimate for the purposes of estimating mutual information and synergy. The decoders we compared were an optimal global linear (GL) decoder, the standard population vector (PV) decoder, and four maximum likelihood decoders each of which differed in terms of their knowledge of the statistics of the stimulus conditioned neural response. For example, consider a global linear decoder

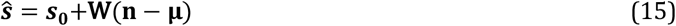

which converts deviations of the neural response, **n**, from the average response over all stimulus values, **μ**, into an estimate via a linear transformation **W.** The linear transformation that minimizes squared error of the estimates takes the form

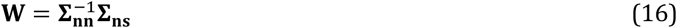

where **∑_nn_** is the total covariance of the neural response (including both stimulus-induced and noise-induced correlations) and **∑_ns_** is the covariance between the neural response and the vector stimulus. This decoder also knows about the mean neural response and mean value of the stimulus vector, but has no understanding of the shape of tuning curves.

Another obvious drawback of the global linear decoder is that it is incapable of accounting for the fact that direction is a circular stimulus variable. The population vector decoder overcomes this deficit but at a cost. For a stimulus vector that represents a circular random variable and a scalar random variable, the population vector decoder actually consists of two independent non-linear decoders, one for each stimulus dimension. The PV operates by determining the preferred speed, 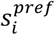, and direction, 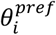, for each neuron. For direction the PV decoder takes a unit vector that points in the preferred direction of a given neuron and then weights that vector by the activity of that neuron. The population vector is a simply a sum of these activity reweighted unit vectors and the decoded direction is simply the direction of the population vector.

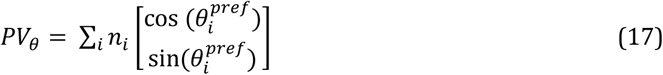

For a scalar variable, the equivalent to a PV is a center of mass decoder. This decoder simply performs a weighted average of preferred stimulus with weight’s given by the neural activity. Defining the population vector as

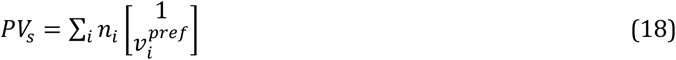

it is easy to show that the tangent of the angle of PV_s_ finds the center of mass. Besides the correct non-linearity, the only things the PV needs to know are the preferred speeds and directions. By analogy to the global linear decoder, we say that the population vector decoder assumes unimodal tuning (a property of the stimulus dependent mean) and ‘knows’ the value of the preferred speed and/or direction of each neuron.

The principle deficiency of the PV’s decoder is that it treats all neurons with the same peak tuning as equally informative despite huge difference in SNR across units. This deficiency is usually demonstrated via comparison to the maximum likelihood estimate^48,49^. For this reason, we also considered the properties of maximum likelihood decoding by computing linear Fisher information, a quantity that can be related to the variance of the locally optimal linear estimator (LOLE). This is equivalent to assuming that the likelihood function takes the form:

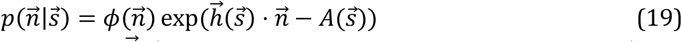

where 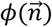 is a largely inconsequential measure function and 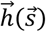 is a stimulus dependent kernel. When 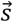 is a continuous valued stimulus such as speed or direction it is easy to show that the gradient of the stimulus dependent kernel is proportional to the weights of a locally optimal linear estimator and thus computable from knowledge of the first and second order statistics of the stimulus conditioned neural response, i.e.

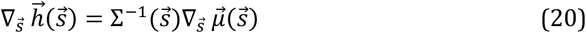

where 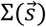 is the stimulus dependent covariance matrix covariance matrix and 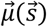 is the stimulus dependent mean. Note the similarity to the globally linear (GL) decoder which depended upon (1) total covariation of the neural response and (2) the covariance between the neural response and the stimulus. Here the stimulus dependent kernel is solely a function of (1) the (possibly stimulus dependent) noise covariance of the neural response and (2) the gradient of the mean response with respect to the stimulus. Indeed, when 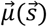 is linear in 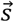 then 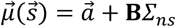 and the proportional relationship between the weights of the LOLE and the gradient of the stimulus-dependent kernel 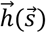 becomes clear.

Because of this relationship between ‘statistical knowledge’ and decoder, we can characterize suboptimal decoders by the statistical properties of stimulus conditioned neural responses that they know, don’t know, or are simply incorrect about. For example, consider a maximum likelihood decoder for direction that is ignorant about speed. A marginal decoder such as this would have knowledge of Σ(*θ*) and 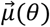, as opposed to knowledge of both the speed and direction dependence of the mean and covariance, i.e. Σ(*θ, ν*) and 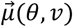. This limited knowledge has the potential to introduce biases and errors in direction estimates that are dependent on speed that are not present when full knowledge of the speed and direction dependence is used to build the decoder. We could also consider maximum likelihood decoders that understand 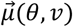 but are limited to knowledge of only the diagonal elements of Σ(*θ, ν*). This is a naïve Bayesian decoder of the joint stimulus (i.e. NB-2D). Similarly, a decoder for direction that was ignorant of speed *and* ignorant of the noise correlations in the neural response (but not the variance) could be called a marginal naïve Bayesian decoder for *θ* (NB-1D). Using this framework, we summarized the decoders tested and their various ‘understandings’ of the statistics of the stimulus conditioned neural response in Table 1.

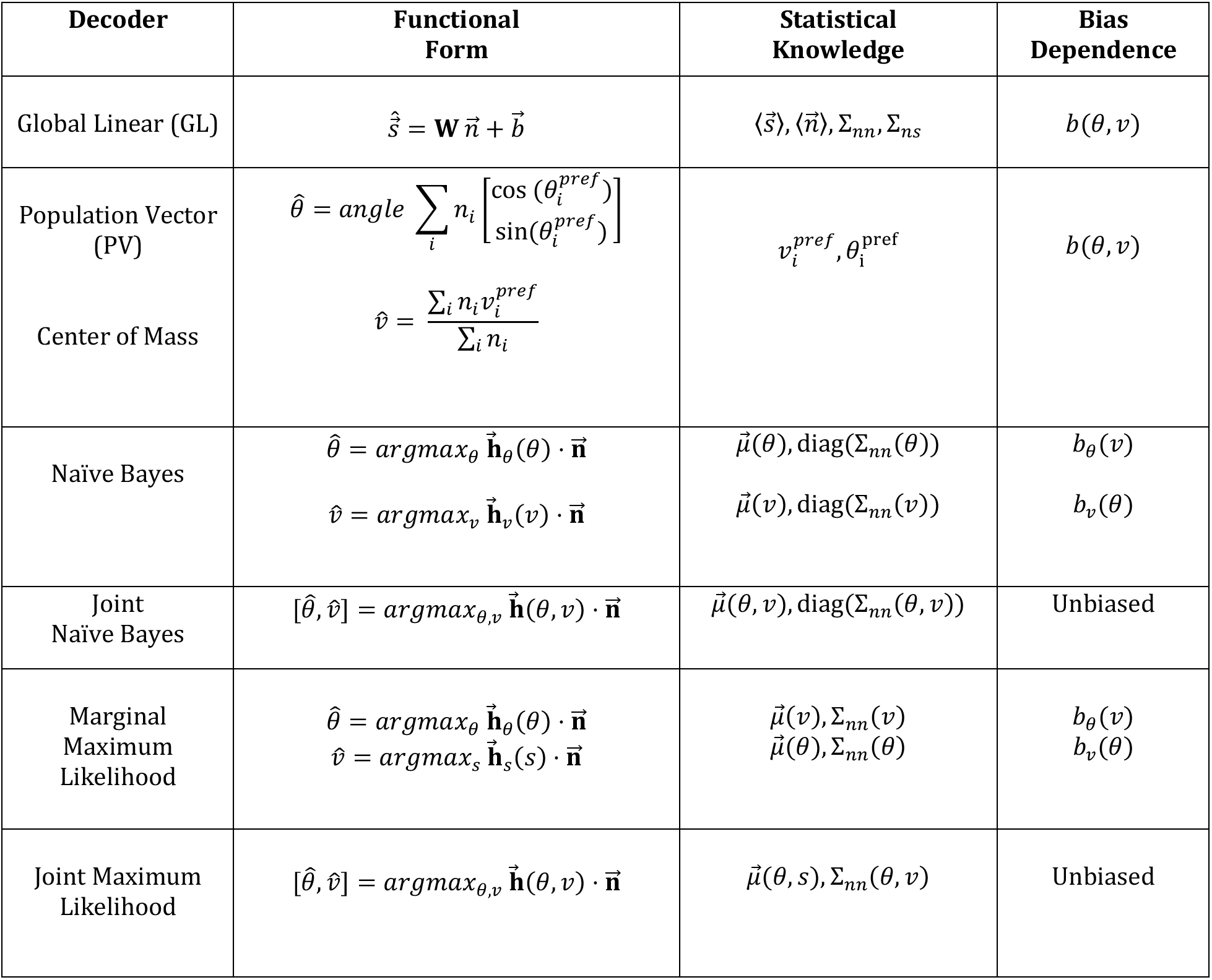

**Table 1.**
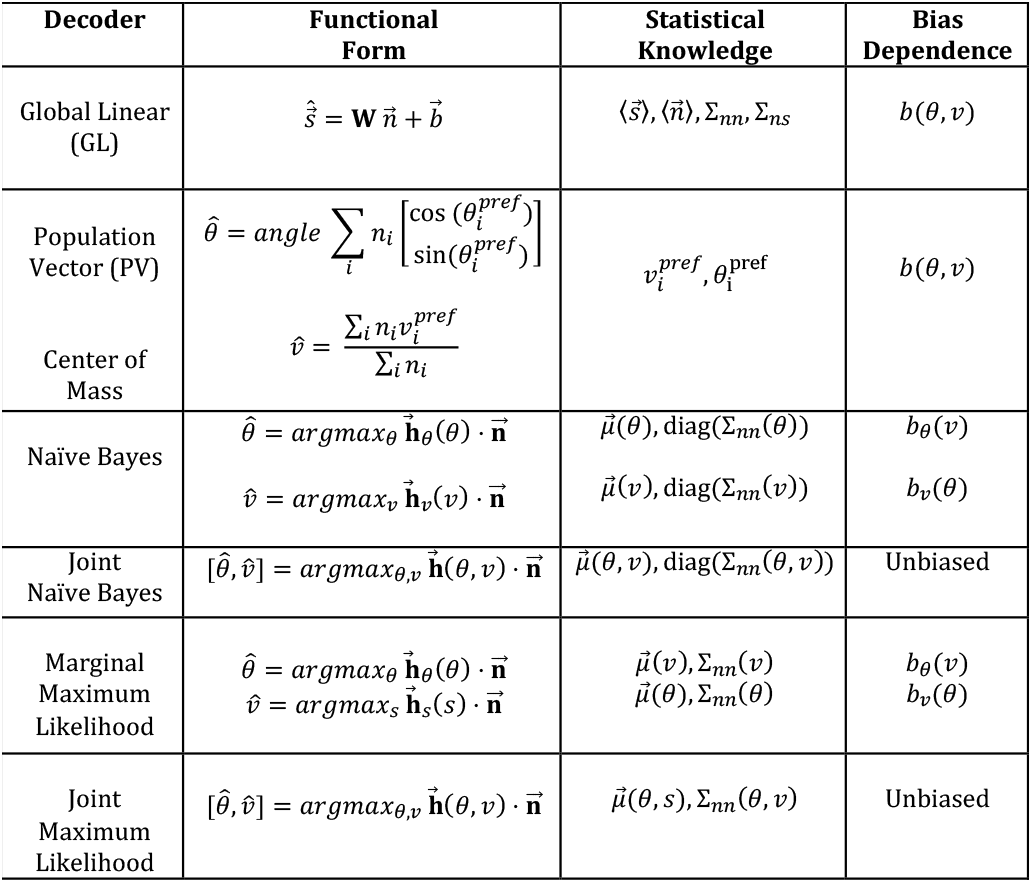

### Stimulus synergy beyond 2D

We extended our analysis of stimulus synergy in single units out to 8 dimensions using a Poisson model with identical tuning along each dimension. To generate Figure 6, we used a normalized stimulus range (in arbitrary units) of −0.5 to 0.5 around the preferred feature for all dimensions. The firing rate was the outer product of all 1-dimensional Gaussian tuning curves:

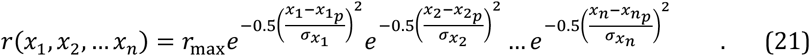

We varied the tuning bandwidth σ of dimensions from 4% to 50% of the arbitrary stimulus range to analyze the impact of tuning sharpness on synergy.

## References

1. Brunel, N. & Nadal, J.-P. P. Mutual Information, Fisher Information, and Population Coding. 10, (1998).

2. Eurich, C. W. & Wilke, S. D. Multidimensional encoding strategy of spiking neurons. Neural Comput. 12, 1519–1529 (2000).

3. Zohary, E. Population coding of visual stimuli by cortical neurons tuned to more than one dimension. Biol. Cybern. 66, 265–272 (1992).

4. Panzeri, S., Treves, A., Schultz, S. & Rolls, E. T. On Decoding the Responses of a Population of Neurons from Short Time Windows. Neural Comput. 11, 1553–1577 (1999).

5. Wilke, S. D. & Eurich, C. W. Representational accuracy of stochastic neural populations. Neural Comput. 14, 155–189 (2002).

6. Montemurro, M. A. & Panzeri, S. Optimal Tuning Widths in Population Coding of Periodic Variables. Neural Comput. 18, 1555–1576 (2006).

7. Benichoux, V., Brown, A. D., Anbuhl, K. L. & Tollin, D. J. Representation of Multidimensional Stimuli: Quantifying the Most Informative Stimulus Dimension from Neural Responses. J. Neurosci. 37, 7332–7346 (2017).

8. Abbott, L. F. & Dayan, P. The effect of correlated variability on the accuracy of a population code. Neural Comput. 11, 91–101 (1999).

9. Ecker, A. S., Berens, P., Tolias, A. S. & Bethge, M. The Effect of Noise Correlations in Populations of Diversely Tuned Neurons. J. Neurosci. 31, 14272–14283 (2011).

10. Brenner, N., Strong, S. P., Koberle, R., Bialek, W. & de Ruyter van Steveninck, R. R. Synergy in a neural code. Neural Comput. 12, 1531–1552 (2000).

11. Reich, D. S., Mechler, F. & Victor, J. D. Independent and Redundant Information in Nearby Cortical Neurons. Science (80-.). 294, (2001).

12. Schneidman, E., Bialek, W. & Berry, M. J. Synergy, redundancy, and independence in population codes. J. Neurosci. 23, 11539–53 (2003).

13. Osborne, L. C., Palmer, S. E., Lisberger, S. G. & Bialek, W. The Neural Basis for Combinatorial Coding in a Cortical Population Response. J. Neurosci. 28, 13522–13531 (2008).

14. Maunsell, J. H. R. & Van Essen, D. C. Functional properties of neurons in middle temporal visual area of the macaque monkey. I. Selectivity for stimulus direction, speed, and orientation. 49, (1983).

15. DeAngelis, G. C. & Uka, T. Coding of Horizontal Disparity and Velocity by MT Neurons in the Alert Macaque. J. Neurophysiol. 89, 1094–1111 (2003).

16. Priebe, N. J., Cassanello, C. R. & Lisberger, S. G. The neural representation of speed in macaque area MT/V5. J. Neurosci. 23, 5650–61 (2003).

17. Newsome, W. T. & Paré, E. B. A selective impairment of motion perception following lesions of the middle temporal visual area (MT). J. Neurosci. 8, 2201–11 (1988).

18. Groh, J. M., Born, R. T. & Newsome, W. T. How is a sensory map read Out? Effects of microstimulation in visual area MT on saccades and smooth pursuit eye movements. J. Neurosci. 17, 4312–4330 (1997).

19. Osborne, L. C., Lisberger, S. G. & Bialek, W. A sensory source for motor variation. Nature (2005). doi:10.1038/nature03961

20. Osborne, L. C., Hohl, S. S., Bialek, W. & Lisberger, S. G. Time Course of Precision in Smooth-Pursuit Eye Movements of Monkeys. J. Neurosci. 27, 2987–98 (2007).

21. Mukherjee, T., Battifarano, M., Simoncini, C. & Osborne, L. C. Shared Sensory Estimates for Human Motion Perception and Pursuit Eye Movements. J. Neurosci. 35, 8515–8530 (2015).

22. Stone, L. S. & Krauzlis, R. J. Shared motion signals for human perceptual decisions and oculomotor actions. J. Vis. 3, 7 (2003).

23. Medina, J. F. & Lisberger, S. G. Variation, signal, and noise in cerebellar sensory-motor processing for smooth-pursuit eye movements. J. Neurosci. 27, 6832–42 (2007).

24. Hohl, S. S. S., Chaisanguanthum, K. S. S. & Lisberger, S. G. G. Sensory population decoding for visually guided movements. Neuron 79, (2013).

25. Chang, L. & Tsao, D. Y. The Code for Facial Identity in the Primate Brain. Cell (2017). doi:10.1016/j.cell.2017.05.011

26. Strong, S. P., Koberle, R., de Ruyter van Steveninck, R. R. & Bialek, W. Entropy and Information in Neural Spike Trains. Phys. Rev. Lett. 80, 197–200 (1998).

27. Osborne, L. C., Bialek, W. & Lisberger, S. G. Time course of information about motion direction in visual area MT of macaque monkeys. J. Neurosci. 24, 3210–22 (2004).

28. Fairhall, A. L. et al. Selectivity for multiple stimulus features in retinal ganglion cells. J. Neurophysiol. (2006). doi:10.1152/jn.00995.2005

29. Mazer, J. A., Vinje, W. E., McDermott, J., Schiller, P. H. & Gallant, J. L. Spatial frequency and orientation tuning dynamics in area V1. Proc. Natl. Acad. Sci. 99, 1645–1650 (2002).

30. Cohen, M. R. & Kohn, A. Measuring and interpreting neuronal correlations. Nat. Neurosci. 14, 811–819 (2011).

31. Ponce-Alvarez, A., Thiele, A., Albright, T. D., Stoner, G. R. & Deco, G. Stimulus-dependent variability and noise correlations in cortical MT neurons. Proc. Natl. Acad. Sci. 110, 13162–13167 (2013).

32. Moreno-Bote, R. R. et al. Information-limiting correlations. 17, 1410–1417 (2014).

33. Sanada, T. M. & DeAngelis, G. C. Neural Representation of Motion-In-Depth in Area MT. J. Neurosci. 34, 15508–15521 (2014).

34. Dayan, P. & Abbott, L. F. Theoretical Neuroscience: Computational and Mathematical Modeling of Neural Systems. Computational and Mathematical Modeling of Neural… (2001).

35. Clark, D. A. et al. Flies and humans share a motion estimation strategy that exploits natural scene statistics. Nat. Neurosci. (2014). doi:10.1038/nn.3600

36. Liu, B., Macellaio, M. V. & Osborne, L. C. Efficient sensory cortical coding optimizes pursuit eye movements. Nat. Commun. 7, 12759 (2016).

37. Grunewald, A. & Skoumbourdis, E. K. The Integration of Multiple Stimulus Features by V1 Neurons. J. Neurosci. 24, 9185–9194 (2004).

38. Angelaki, D. E., Gu, Y. & DeAngelis, G. C. Multisensory integration: psychophysics, neurophysiology, and computation. Curr. Opin. Neurobiol. 19, 452–458 (2009).

39. Gu, Y., Cheng, Z., Yang, L., DeAngelis, G. C. & Angelaki, D. E. Multisensory Convergence of Visual and Vestibular Heading Cues in the Pursuit Area of the Frontal Eye Field. Cereb. Cortex 26, 3785–3801 (2016).

40. Schneidman, E. et al. Synergy from Silence in a Combinatorial Neural Code. J. Neurosci. 31, 15732–15741 (2011).

41. Heeger, D. J., Simoncelli, E. P. & Movshon, J. A. Computational models of cortical visual processing. Proc. Natl. Acad. Sci. U. S. A. 93, 623–7 (1996).

42. Beck, J. M., Latham, P. E. & Pouget, A. Marginalization in Neural Circuits with Divisive Normalization. J. Neurosci. (2011). doi:10.1523/JNEUROSCI.1706-11.2011

43. Lisberger, S. G. & Movshon, J. A. Visual motion analysis for pursuit eye movements in area MT of Macaque monkeys. J. Neurosci. 19, 2224–2246 (1999).

44. DeAngelis, G. C., Cumming, B. G. & Newsome, W. T. Cortical area MT and the perception of stereoscopic depth. Nature (1998). doi:10.1038/29299

45. Lisberger, S. G. & Westbrook, L. E. Properties of visual inputs that initiate horizontal smooth pursuit eye movements in monkeys. J. Neurosci. 5, 1662–73 (1985).

46. Xu, X., Collins, C. E., Khaytin, I., Kaas, J. H. & Casagrande, V. A. Unequal representation of cardinal vs. oblique orientations in the middle temporal visual area. Proc. Natl. Acad. Sci. (2006). doi:10.1073/pnas.0608502103

47. Priebe, N. J., Lisberger, S. G. & Movshon, J. A. Tuning for spatiotemporal frequency and speed in directionally selective neurons of macaque striate cortex. J. Neurosci. 26, 2941–50 (2006).

48. Series, P., Latham, P. E. & Pouget, A. Tuning curve sharpening for orientation selectivity: Coding efficiency and the impact of correlations. Nat. Neurosci. (2004). doi:10.1038/nn1321

49. Sharpee, T. O. & Berkowitz, J. A. Linking neural responses to behavior with informationpreserving population vectors. Current Opinion in Behavioral Sciences (2019). doi:10.1016/j.cobeha.2019.03.004

